# The WD and linker domains of ATG16L1 required for non-canonical autophagy limit lethal respiratory infection by influenza A virus at epithelial surfaces

**DOI:** 10.1101/2020.01.15.907873

**Authors:** Yingxue Wang, Weijiao Zhang, Matthew Jefferson, Parul Sharma, Ben Bone, Anja Kipar, Janine L. Coombes, Timothy Pearson, Angela Man, Alex Zhekova, Yongping Bao, Ralph A Tripp, Yohei Yamauchi, Simon R. Carding, Ulrike Mayer, Penny P. Powell, James P. Stewart, Thomas Wileman

## Abstract

Respiratory viruses such as influenza A virus (IAV) and SARS-CoV-2 (Covid-19) cause pandemic infections where cytokine storm syndrome, lung inflammation and pneumonia lead to high mortality. Given the high social and economic cost of these viruses, there is an urgent need for a comprehensive understanding of how the airways defend against virus infection. Viruses entering cells by endocytosis are killed when delivered to lysosomes for degradation. Lysosome delivery is facilitated by non-canonical autophagy pathways that conjugate LC3 to endo-lysosome compartments to enhance lysosome fusion. Here we use mice lacking the WD and linker domains of ATG16L1 to demonstrate that non-canonical autophagy protects mice from lethal IAV infection of the airways. Mice with systemic loss of non-canonical autophagy are exquisitely sensitive to low-pathogenicity murine-adapted IAV where extensive viral replication throughout the lungs, coupled with cytokine amplification mediated by plasmacytoid dendritic cells, leads to fulminant pneumonia, lung inflammation and high mortality. IAV infection was controlled within epithelial barriers where non-canonical autophagy slowed fusion of IAV with endosomes and reduced activation of interferon signalling. This was consistent with conditional mouse models and *ex vivo* analysis showing that protection against IAV infection of lung was independent of phagocytes and other leukocytes. This establishes non-canonical autophagy pathways in airway epithelial cells as a novel innate defence mechanism that can restrict IAV infection and lethal inflammation at respiratory surfaces.

## Introduction

Influenza A virus (IAV) is a respiratory pathogen of major global public health concern (Yamayoshi & Kawaoka, 2019). As with SARS CoV-2, animal reservoirs of IAV can contribute to zoonotic infection leading to pandemics with a high incidence of viral pneumonia, morbidity and mortality. IAV infects airway and alveolar epithelium and damage results from a combination of the intrinsic pathogenicity of individual virus strains as well as the strength and timing of the host innate/inflammatory responses. Optimal cytokine levels protect from IAV replication and disease but excessive cytokine production and inflammation worsens the severity of lung injury (Davidson et al., 2014, Herold et al., 2015, Iwasaki & Pillai, 2014, Ramos & Fernandez-Sesma, 2015, Teijaro et al., 2014). Even though infection of the lower respiratory tract can result in inflammation, flooding of alveolar spaces, acute respiratory distress syndrome and respiratory failure the factors that control IAV replication at epithelial surfaces and limit lethal lung inflammation remain largely unknown.

The transport of viruses to lysosomes for degradation provides an important barrier against infection. Transport to lysosomes can be enhanced non-canonical autophagy pathways which conjugate autophagy marker protein LC3 to endo-lysosome compartments to increase lysosome fusion. In phagocytes LC3-associated phagocytosis (LAP) conjugates LC3 to phagosomes and enhances phagosome maturation (Delgado et al., 2008, Fletcher et al., 2018, Lamprinaki et al., 2017, Martinez et al., 2015, Sanjuan et al., 2007). In non-phagocytic cells LC3 is conjugated to endo-lysosome compartments during the uptake of particulate material such as apoptotic cells and aggregated β-amyloid, and following membrane damage during pathogen entry or osmotic imbalance induced by lysosomotropic drugs (Heckmann et al., 2019, Tan et al., 2018) (Florey et al., 2015, Florey et al., 2011, Roberts et al., 2013). It is known from *in vitro* studies that LC3 can be recruited to endo-lysosome compartments during the uptake of pathogens, but the roles played by non-canonical autophagy during viral infection *in vivo* are largely unknown.

A role for non-canonical autophagy in host defence has been implied from *in vitro* studies of LAP in phagocytes infected with free living microbes with a tropism for macrophages such as bacteria (*Listeria monocytogenes* (Gluschko et al., 2018), *Legionella dumoffii* (Hubber et al., 2017)), protozoa (*Leishmania major*) and fungi (*Aspergillus fumigatus* (Akoumianaki et al., 2016, Kyrmizi et al., 2018, Matte et al., 2016)). It is also known that IAV induces non-canonical autophagy during infection of cells in culture (Fletcher et al., 2018), however, the role played by non-canonical autophagy in controlling IAV infection and lung inflammation *in vivo* are currently unknown. It is not known for example if non-canonical autophagy is important in the control of IAV infection by epithelial cells at sites of infection, or if it plays a predominant role within phagocytes and antigen-presenting cells during development of an immune response. Herein we use mice with specific loss of non-canonical autophagy to determine the role played by non-canonical autophagy in host defence against IAV infection of the respiratory tract. The mice (δWD) lack the WD and linker domains of ATG16L1 that are required for conjugation of LC3 to endo-lysosome membranes (Rai et al., 2019) but express the N-terminal ATG5-binding domain and the CCD of ATG16L1 that are required for WIPI2 binding and autophagy (Dooley et al., 2014). Importantly, the δWD mice grow normally and maintain tissue homeostasis (Rai et al., 2019), and unlike autophagy-defective mice, the δWD mice do not have pro-inflammatory phenotype.

We show that loss of non-canonical autophagy from all tissues renders mice highly sensitive to low-pathogenicity murine-adapted IAV (A/X-31) leading to extensive viral replication throughout the lungs, cytokine dysregulation and high mortality typically seen after infection with highly pathogenic IAV. Conditional mouse models and *ex vivo* analysis showed that protection against IAV infection of lung was independent of phagocytes and other leukocytes, and that infection was controlled within epithelial barriers where non-canonical autophagy slowed fusion of IAV with endosomes and reduced interferon signalling. This establishes non-canonical autophagy pathways in airway epithelial cells as a novel innate defence mechanism that restricts IAV infection at respiratory surfaces.

## Results

### Mice with systemic loss of the WD and linker domains of ATG16L1 are highly sensitive to IAV infection

The consequences of loss of the WD and linker domains of ATG16L1 on conventional autophagy and non-canonical autophagy were confirmed using cell lines taken from controls and δWD mice. Mouse embryo fibroblasts (MEFs) from littermate control mice expressed full-length α and β forms of ATG16L1 at 70kDa (Fig. 1B), and generated PE-conjugated LC3II during recruitment of LC3 to autophagosomes following starvation in HBSS. The MEFs also recruited LC3 to endo-lysosome compartments swollen by monensin and control bone marrow-derived macrophages (BMDM) activated LAP to recruit LC3 to phagosomes containing zymosan (Fig. 1B). MEFs from δWD mice expressed a truncated ATG16L1 at 30 kDa (Fig. 1C). Cells from δWD mice generated LC3II and autophagosomes in response to starvation but failed to recruit LC3 to swollen endo-lysosome compartments or phagosomes containing zymosan. These data confirm defects in non-canonical autophagy and LAP in the δWD mice.

**Fig. 1.**
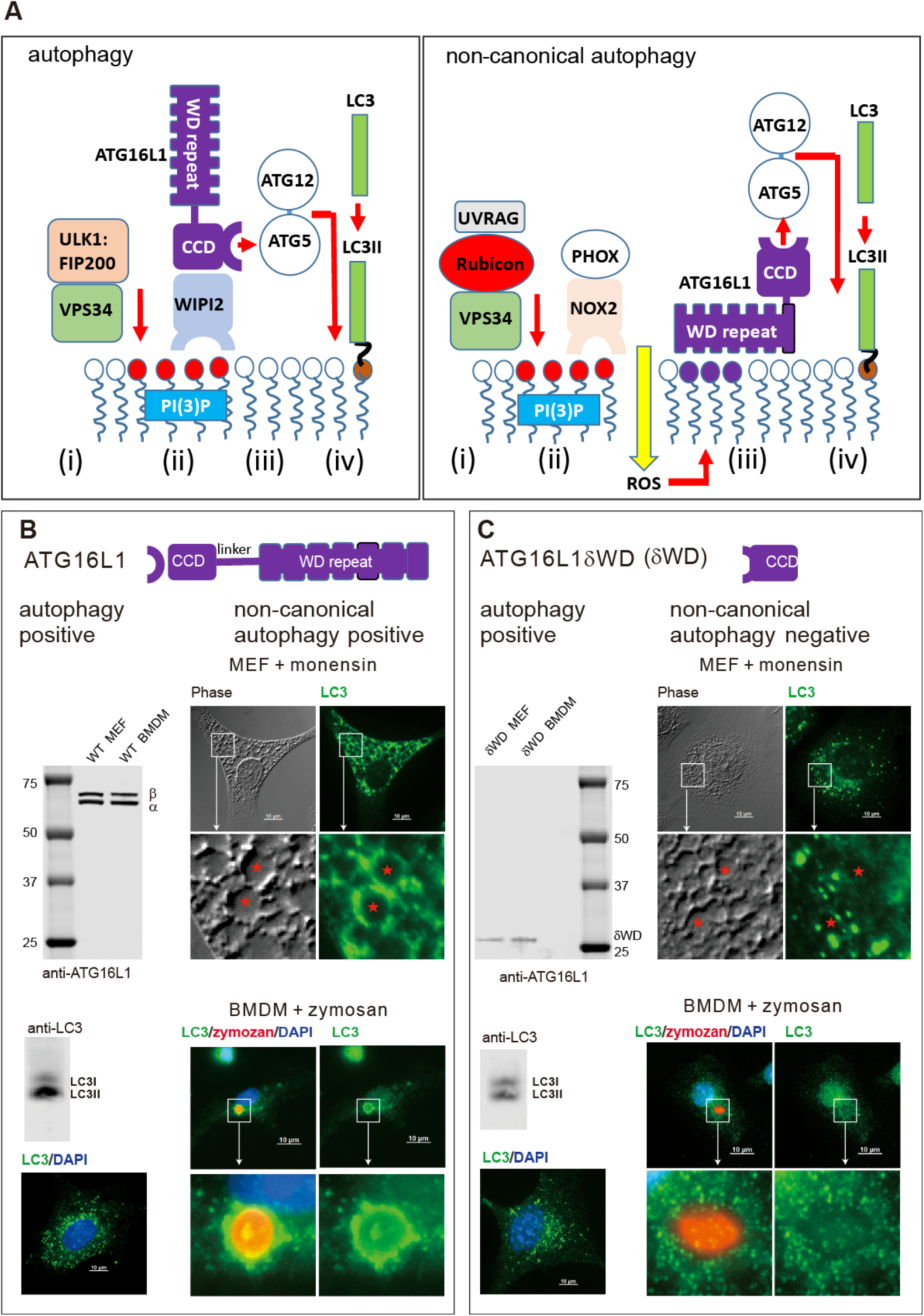
Deletion of the WD domain of ATG16L1 causes loss of non-canonical autophagy and LAP but retains autophagy. **A)**. Conventional autophagy is activated by the initiation complex containing ULK1, FIP200 and the PI3 kinase VPS34 (i) which responds to starvation. The VPS34 subunit phosphorylates lipids in membranes to generate sites for WIP2 binding (ii). WIPI2 binds the coiled coil domain (CCD) of ATG16L1 (ii) leading to recruitment of ATG16L1 and the LC3 conjugation machinery (ATG16l1:ATG5-ATG12, ATG7, ATG3) (iii). This results in conjugation of LC3 to PE in the surface of the autophagosome (iv) to promote fusion with lysosomes. During non-canonical autophagy signalling pathways arising from the lumen of the endosome or phagosome recruit a complex containing UVRAG, Rubicon and VPS34 (i). The VPS34 subunit phosphorylates lipids in endo-lysosome membranes to generate sites for binding the multicomponent PHOX:NOX2 complex (ii) which is stabilised by Rubicon to generate reactive oxygen species (ROS). ROS induces binding of the WD domain of ATG16L1 to endo-lysosome membranes (iii) leading to recruitment of the LC3 conjugation machinery (ATG16l1:ATG5-ATG12, ATG7, ATG3) (iii) and conjugation of LC3 to PE (iv) **B) Left column:** MEFs from littermate control express α and β isoforms of ATG16L1 at 70 kDa, convert LC3I to LC3II and generate LC3 puncta during autophagy induced by HBSS. **Right column:** LC3 (green) is recruited to endo-lysosomes following induction of non-canonical autophagy by monensin, and to phagosomes following engulfment of zymosan by bone marrow-derived macrophages (BMDM). **C) Left column:** MEFs from δWD mice express a 30kDa truncated ATG16L1, but still convert LC3I to LC3II and generate LC3 puncta following autophagy induced by HBSS. **Right column:** The δWD MEFs do not recruit LC3 (green) to endo-lysosomes following incubation with monensin or to bone marrow-derived macrophage phagosomes containing zymosan.

IAV enters airway and lung epithelial cells by endocytosis, and in tissue culture IAV induces non-canonical autophagy leading to ATG16L1-WD domain-dependent conjugation of LC3 to the plasma membrane and peri-nuclear structures (Fletcher et al., 2018). To test whether non-canonical autophagy has a host defence function *in vivo*, δWD mice were infected with IAV. We used a low-pathogenicity murine-adapted IAV (A/X31) that does not normally lead to extensive viral replication throughout the lungs, or cause the cytokine storm syndrome and death typically seen after infection with highly pathogenic viral strains. The results (Fig. 2) showed that δWD mice became moribund and showed severe signs of clinical illness (rapid breathing, piloerection). They also displayed rapid weight loss compared to littermate controls (Fig. 2A) and had increased mortality with survivors recovering more slowly from infection (Fig. 2B). The increased weight loss was associated with an approx. log increase in lung virus titre at 5 days post-infection (d.p.i.; Fig. 2C). Furthermore, histopathology and immunohistochemistry (IH) analysis of lungs from δWD mice showed fulminant viral pneumonia with large numbers of IAV-positive cells (Fig. 2D).

**Fig. 2.**
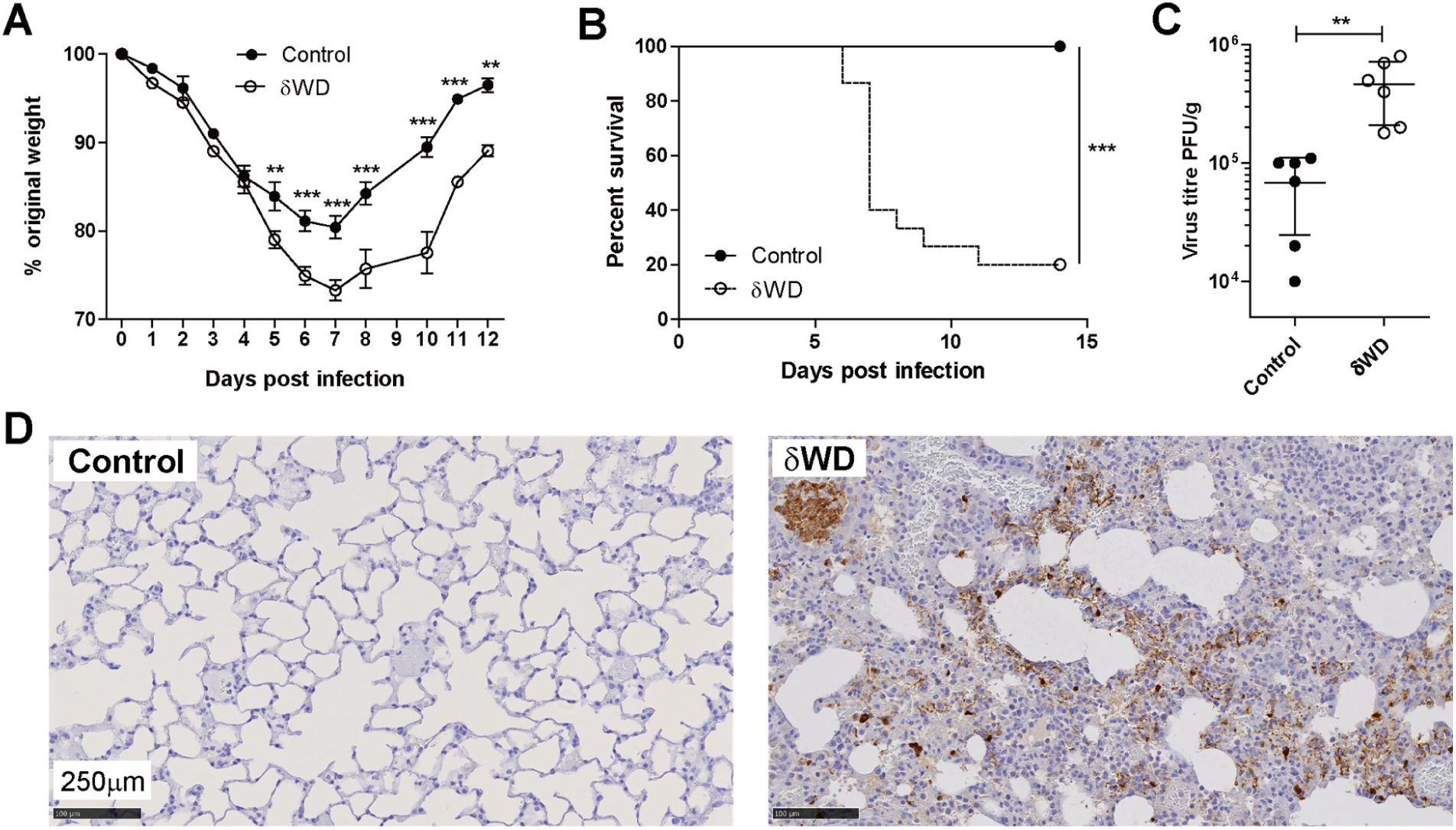
Systemic loss of non-canonical autophagy increases susceptibility to IAV infection. Littermate control and δWD mice were challenged intranasally with IAV strain X31 (10^3^ pfu). **(A)** Mice were monitored for weight loss at indicated time-points. (n = 8). Data represent the mean value ± SEM. Comparisons were made using a repeated-measures two-way ANOVA (Bonferroni post-test). **(B)** Survival was assessed at indicated time points (n = 15). Comparisons were made using log-rank (Mantel-Cox) test **(C)** IAV titre in lungs was determined by plaque assay at 5 d.p.i. (n = 6). Data for individual animals are shown, bars represent the mean ± SD. Mann-Whitney *U* test was used to determine significance. **(D)** The presence of IAV antigen was assessed by IH at 7 d.p.i. (representative images from n = 6).

### Non-canonical autophagy controls lung inflammation after IAV infection

Innate protection against IAV is provided by type 1 (α, β) and III (λ) interferon (IFN) with severe IAV infection causing excessive airway inflammation and pulmonary pathology attributable in part to IFNαβ and TNF-α (Davidson et al., 2014, Szretter et al., 2007). Measurement of cytokine expression at 2 d.p.i showed that IAV induced a transient increase in transcripts for interferon-stimulated genes (ISGs), ISG15 and IFIT1 (Iwasaki & Pillai, 2014) and pro-inflammatory cytokines (IL-1β, TNF-α, and CCL2 [MCP-1]) in the lungs of both control and δWD mice (Fig 3A). This increase in cytokine expression was resolved by 3 d.p.i. before a second wave of increased cytokine expression at 5 d.p.i. This second wave of cytokine expression was resolved by 7 d.p.i in control mice, but δWD mice showed sustained increases in ISG15, IFIT1, IL-1β, TNF-α and CCL2 transcripts, co-incident with exacerbated weight loss. At 3 d.p.i lungs of δWD mice showed increased expression of neutrophil chemotaxis factor CXCL1 mRNA (Fig. 3A), coincident with increased neutrophil infiltration of airways and parenchyma, and extensive neutrophil extracellular traps (NETs) as a consequence of neutrophil degeneration as shown by IH (Fig. 3B and S1). Increased neutrophil infiltration of airways in δWD mice at 2 d.p.i. was confirmed and quantified using flow cytometric analysis of broncho-alveolar lavage (BAL; Fig. 3C). At 5 - 7 d.p.i. increased expression of CCL2 mRNA in δWD mice was coincident with extensive macrophage/monocyte infiltration into lung parenchyma observed by IH (Fig. 3B and S2) which was not seen in controls. This increased macrophage/monocyte infiltration in δWD mice was confirmed and quantified using flow cytometric analysis of single cell suspensions from lung tissue (Fig. 3D). It is known that, in severe IAV infection, a cytokine storm occurs that is amplified by plasmacytoid dendritic cells pDCs (Davidson et al., 2014). pDCs detect virus-infected cells and produce large amounts of cytokines, in particular IFNαβ, that in severe infections can enhance disease. In these cases, depletion of pDCs can decrease morbidity (Davidson et al., 2014). Depletion of pDCs in IAV-infected δWD mice using anti-PDCA-1 led to markedly decreased weight loss as compared with isotype control-treated mice and that was similar to that seen in littermate controls (Fig3E). This indicates that excessive cytokine production amplified by pDCs is responsible for the increased morbidity seen in the δWD mice.

**Fig. 3.**
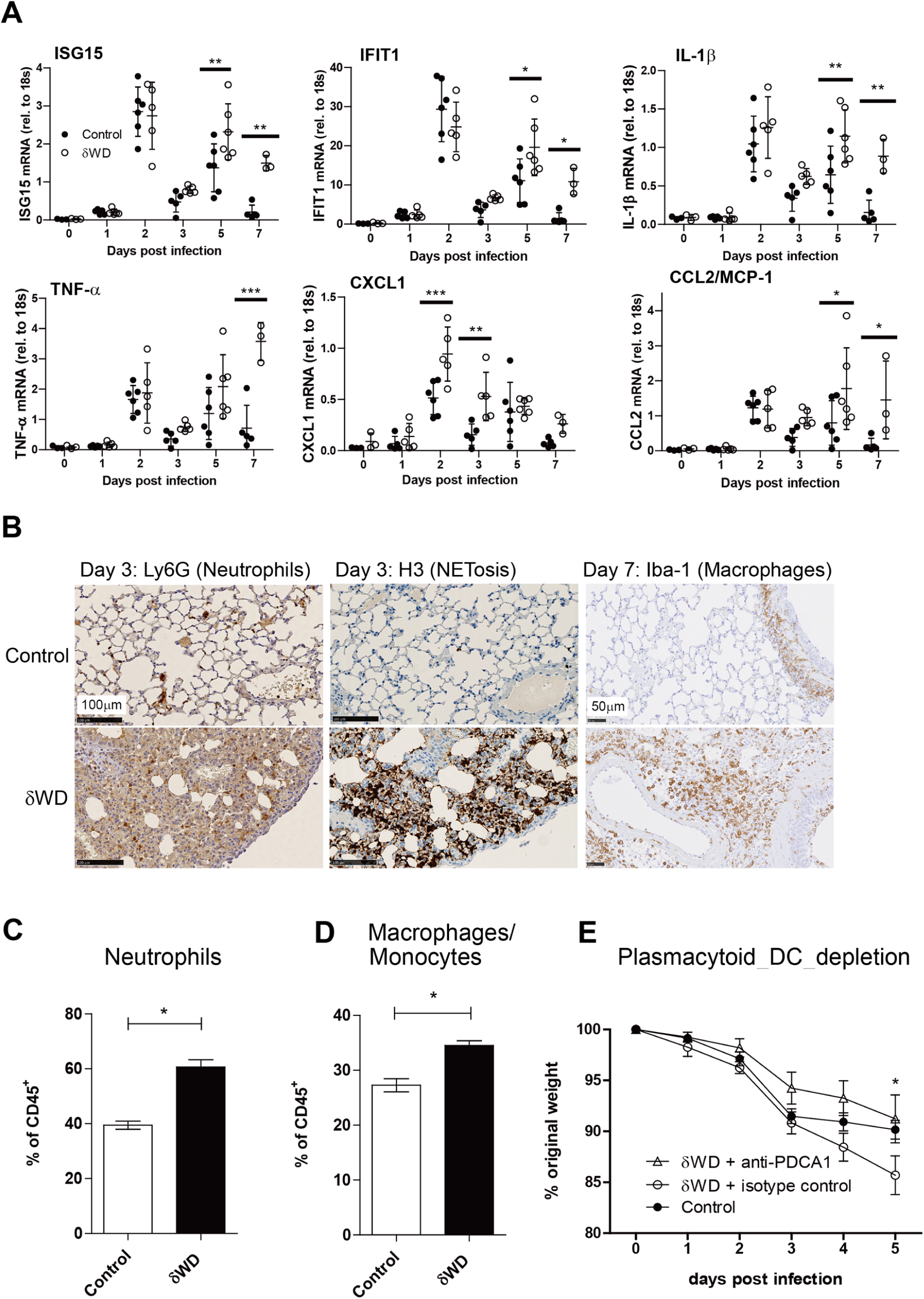
Systemic loss of non-canonical autophagy leads to extensive lung inflammation and damage. Littermate control and δWD mice (n = 5) were challenged with IAV X31 (10^3^ pfu). (**A)** At the indicated time points, cytokine mRNA transcripts in lung tissue (n = 5) were evaluated by qPCR. Data for individual animals are shown, bars represent the mean ± SD and were compared by 2-way ANOVA with Bonferroni post-tests. (**B**) Representative lung sections from animals (n = 6) taken at 3 d.p.i. were stained by IH for neutrophils (Ly6G) or neutrophil extracellular traps (NET; anti-H3). Sections at 7 d.p.i. were stained for macrophages (Iba-1). Further micrographs are shown in Figs. S1 and S2. **(C)**. BAL (n = 5) was taken at 2 d.p.i and evaluated by flow cytometry, with pre-gating on CD45^+^. The percentage of neutrophils (CD11b^+^, Ly6G^+^) cells is shown ±SEM and were compared using Mann-Whitney *U* test. **(D)**. Single cell suspensions were prepared from lungs taken at 5 d.p.i and evaluated by flow cytometry, with pre-gating on CD45^+^. The percentage of macrophage/monocytes (CD11b^+^, F4/80^+^) cells is shown ±SEM and were compared using Mann-Whitney *U* test. **(E)**. δWD mice were treated with either anti-PCDA-1 (to deplete plasmacytoid DC) or an isotype-matched control. Litter-mate control mice were used as comparator. Weight loss was measured at the indicated days p.i. (n = 5). Comparisons were made using a repeated-measures two-way ANOVA (Bonferroni post-test)

Thus, mice with systemic loss of non-canonical autophagy failed to control lung virus replication and inflammation, leading to increased cytokine production, morbidity and mortality.

### Systemic loss of the WD and linker domains of ATG16L1 does not lead to gross changes in inflammatory threshold or immunological homeostasis

Macrophages cultured from embryonic livers from mice with complete loss of ATG16L1 secrete high levels of IL1-β (Saitoh et al., 2008), and LysMcre-mediated deletion of genes essential for conventional autophagy (eg: Atg5, Atg7, Atg14, Atg16L1, FIP200) in mice leads to raised pro-inflammatory cytokine expression in the lung. This has been reported to increase resistance to IAV infection (Lu et al., 2016), and this was also observed in mice used in our study (Fig. S3) where LysMcre-mediated loss of Atg16L1 prevented rapid weight loss and reduced virus titre. This led us to test the possibility that the δWD mutation to ATG16L1 could also increase IL-1β secretion, and cause the increased inflammation observed during IAV infection. This was tested by incubating BMDM with LPS and purine receptor agonist, BzATP (Fig S4A), or by challenging mice with LPS (Fig S4B). Mice with a complete loss of ATG16L1 in myeloid cells (Atg16L1^fl/fl^-lysMcre) showed three-fold increases in IL-1β in both serum and of secretion IL-1β from BMDM *in vitro*. In contrast IL-1β secretion in δWD mice did not differ significantly from littermate controls (Fig S4A&B). This was consistent with lack of elevated cytokines in lungs prior to infection (see day 0 in Fig. 3A), and our previous work showing that serum levels of IL-1β, IL-12p70, IL-13, and TNF-α in δWD mice are the same as in littermate controls at 8-12 and 20-24 weeks (Rai et al., 2019). The exaggerated inflammatory response to IAV in δWD mice did not therefore result from a raised pro-inflammatory threshold or dysregulated IL-1β responses in the lung. Also, the frequencies of T-cell, B-cell and macrophages were similar in δWD mice to littermate controls (Fig. S5). These data suggest that the exaggerated responses of δWD mice to IAV do not occur because the mice have a raised inflammatory threshold or abnormal immunological homeostasis.

### Non-canonical autophagy limits IAV infection independently of phagocytic cells

The link between non-canonical autophagy/LAP, TLR signalling, NADPH oxidase activation and ROS production (Delgado et al., 2008, Martinez et al., 2015, Sanjuan et al., 2007) provides phagocytes with a powerful mechanism to limit infections *in vivo*. To test whether wild-type bone marrow-derived cells could protect susceptible δWD mice from lethal IAV infection, we generated radiation chimeras (Fig. S6). When challenged with IAV, δWD mice reconstituted with either wild-type or δWD bone marrow remained highly sensitive to IAV (Fig. 4A & B) with body weight reduced by up to 25% and decreased survival by 5 d.p.i. As seen for δWD mice, weight loss was associated with a 10-fold increase in lung viral titre (Fig. 4C), fulminant pneumonia and inflammatory infiltration into the lung (Fig. 4D). This increased susceptibility to IAV was not observed for control mice reconstituted with wild-type marrow, showing that non-canonical autophagy pathways in phagocytes and other leukocytes from control mice were not able to protect δWD mice against lethal IAV infection. In a reciprocal experiment (Fig 5) mice expressing Cre recombinase in myeloid cells (LysMcre) were used to generate mice (called δWD^phag^), where the truncated *Atg16*L1δWD gene was restricted to phagocytic cells (Fig S7). In these mice, non-canonical autophagy was absent in cultured phagocytes (BMDM) but it was present in skin fibroblasts (Fig S7D). After infection with IAV, δWD^phag^ mice showed comparable weight loss and virus titres to those seen in littermate control mice (Fig. 5A & B). Likewise, the raised IL-1β levels (Fig. 5C) and profuse macrophage and neutrophil lung infiltration observed in δWD mice were absent (Fig. S8) and similar to littermate controls. The ability of the WD and linker domains of ATG16L1 to protect epithelial cells against IAV infection was tested *ex vivo* to further exclude any contribution from recruited leukocytes. Virus titres in precision cut lung slices (Fig. 5D) from δWD mice were 10-fold greater than controls. Thus, the sensitivity of δWD mice to IAV was not due to the loss of non-canonical autophagy from myeloid cells, making it likely that non-canonical autophagy mediated by the WD and linker domains of ATG16L1 protects against lethal IAV infection in non-myeloid tissue.

**Fig. 4.**
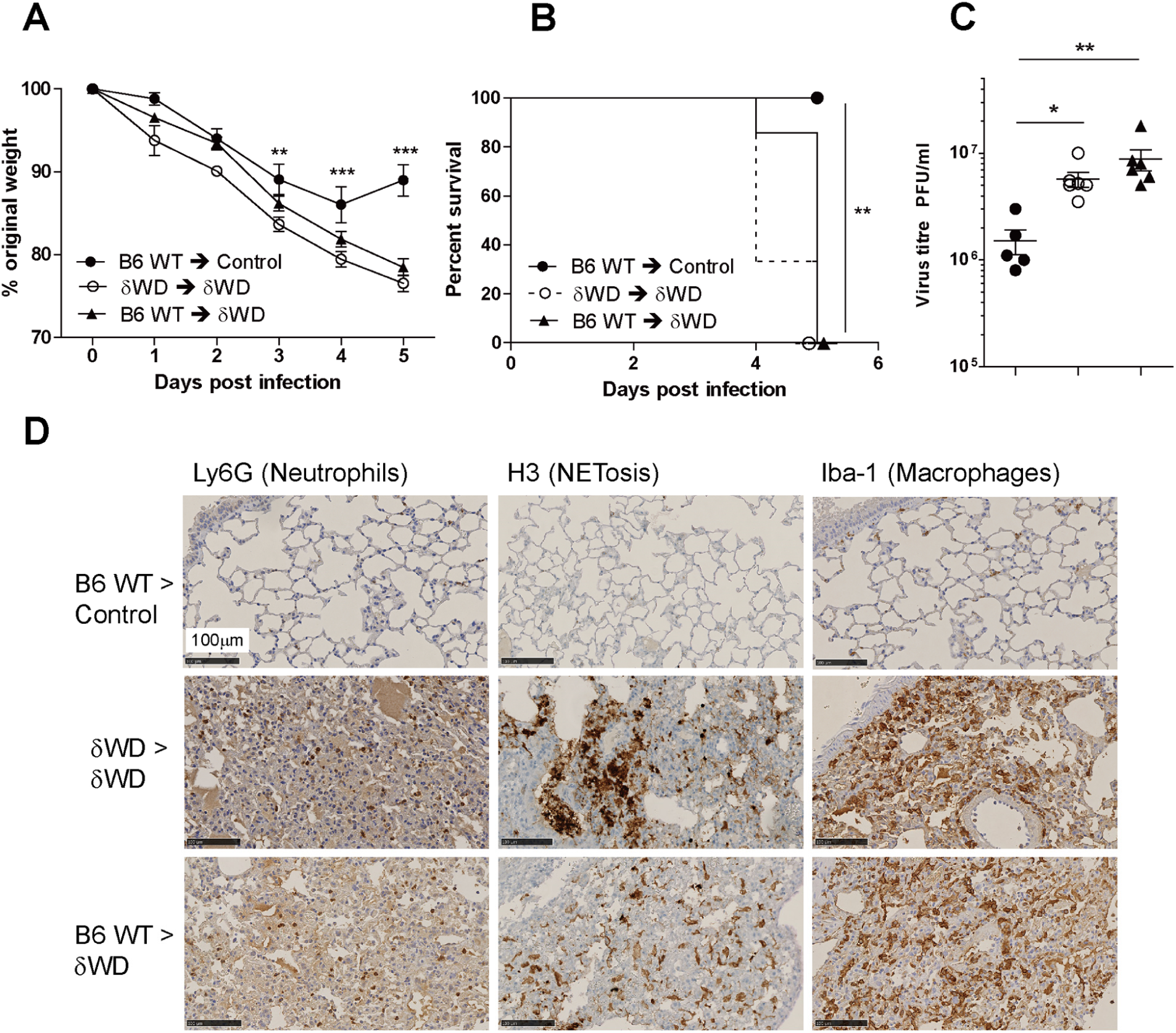
Bone marrow-derived lymphoid tissue cannot reverse sensitivity to IAV infection. Bone marrow from wild-type (*Atg16*L1^+/+^) was used to reconstitute irradiated littermate control mice (B6 WT → control [●]) or δWD mice (B6 WT → δWD [○]). Bone marrow from δWD mice was used to reconstitute irradiated δWD mice (δWD → δWD [▲]). After 12 weeks, mice (n = 5 per group) were challenged with IAV X31 (10^3^ pfu). **(A)** Mice were monitored for weight loss at indicated time-points. Data represent the mean value ± SEM. Comparisons were made using a repeated-measures two-way ANOVA (Bonferroni post-test). **(B)** Survival was assessed at indicated time points. Comparisons were made using log-rank (Mantel-Cox) test **(C)** IAV titre in lungs was determined by plaque assay at 5 d.p.i. (n = 6). Data for individual animals are shown, bars represent the mean ± SD. A one-way ANOVA with Tukey’s post-hoc analysis was used to determine significance. **(D)** Lungs taken at 5 d.p.i. were analysed for neutrophils (Ly6G), neutrophil extracellular traps (NET; anti-H3) and macrophages (Iba-1)

**Fig. 5.**
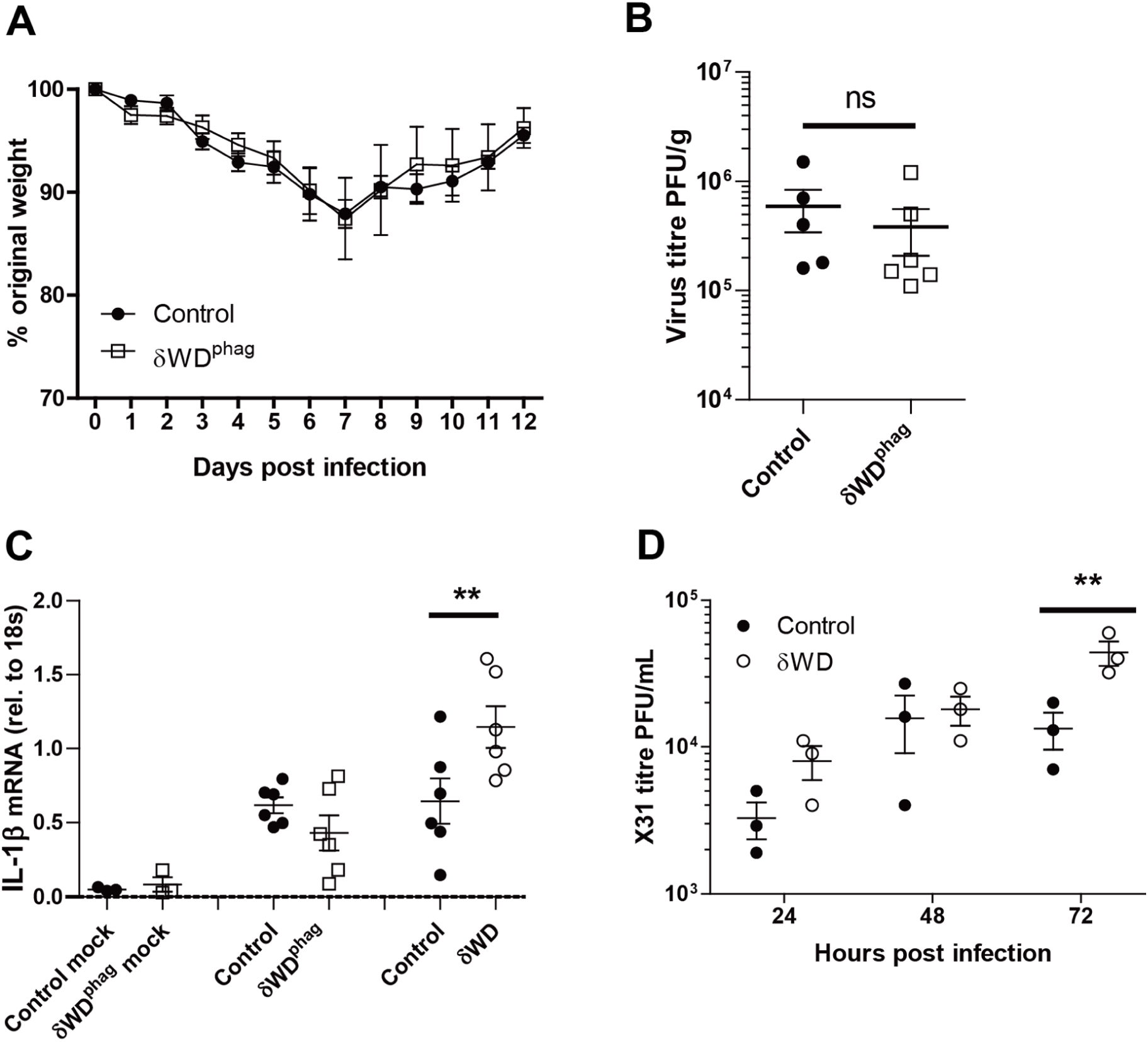
Non canonical autophagy suppresses IAV replication independently of LAP in phagocytes *in vivo* and *in vitro*. Panels **A to C** δWD^phag^ mice lack non-canonical autophagy in myeloid (LysMcre) cells (for construction see Fig. S6A). Offspring negative for LysMcre were used as littermate controls. Mice (n = 6 per group) were challenged intranasally with IAV X31 (10^3^ pfu). **(A)** Mice were monitored for weight loss at indicated time-points. Data represent the mean value ± SEM. Comparisons were made using a repeated-measures two-way ANOVA (Bonferroni post-test). **(B)** IAV titre in lungs was determined by plaque assay at 5 d.p.i. (n = 6). Data for individual animals are shown, bars represent the mean ± SD. Mann-Whitney *U* test was used to determine significance. **(C)** IL-1β mRNA transcripts in lung at 5 d.p.i. were determined by qPCR. Mann-Whitney *U* test was used to determine significance. **(D)** Precision-cut lung slices from control and δWD mice were infected with IAV. Virus titres were determined at indicated time points. Comparisons were made using two-way ANOVA with Bonferroni post-tests.

### Non-canonical autophagy slows IAV fusion with endosomes and reduces interferon signalling

IAV enters cells by receptor-mediated endocytosis where acidification of late endosomes results in fusion with the endosomal membrane and delivery of viral ribonuclear proteins (RNPs) into the cytoplasm (Skehel & Wiley, 2000, Wharton et al., 1994). RNPs are then imported into the nucleus for genome replication (Boulo et al., 2007). The effect of non-canonical autophagy on IAV entry was tested using a fluorescence de-quenching assay where the envelope of purified IAV was labelled with green (DiOC18) and red (R18) lipophilic dyes. MEFs incubated with IAV for increasing times were analyzed by FACs to detect the green fluorescence signal generated when the dyes are diluted following IAV fusion with the endosome membrane (Fig. 6A-C). The percentage of cells emitting a green signal was greater in MEFs from δWD mice (60% compared to 40% for controls at 30 min p.i.) and increased with time (73% versus 56% for controls; Fig. 6B). Likewise, the median fluorescence intensity was 1.6 – 1.4-fold higher in MEFs from δWD mice (Fig 6C). This showed that non-canonical autophagy slowed fusion of IAV with endosome membranes. The recognition of viral RNA by interferon sensors following delivery of RNPs into the cytoplasm was used as a second assay for IAV entry. MEFs from δWD mice showed between 3 and 5-fold increases in expression of IFN responsive genes, ISG15 and IFIT1 (Fig. 6D & E), and this was also observed in the lung *in vivo* (Fig. 3A). Taken together the results demonstrate for the first time that the WD and linker domains of ATG16L1 allow non-canonical autophagy to provide a novel innate defence mechanism against lethal IAV infection within the epithelial barrier *in vivo*.

**Fig. 6.**
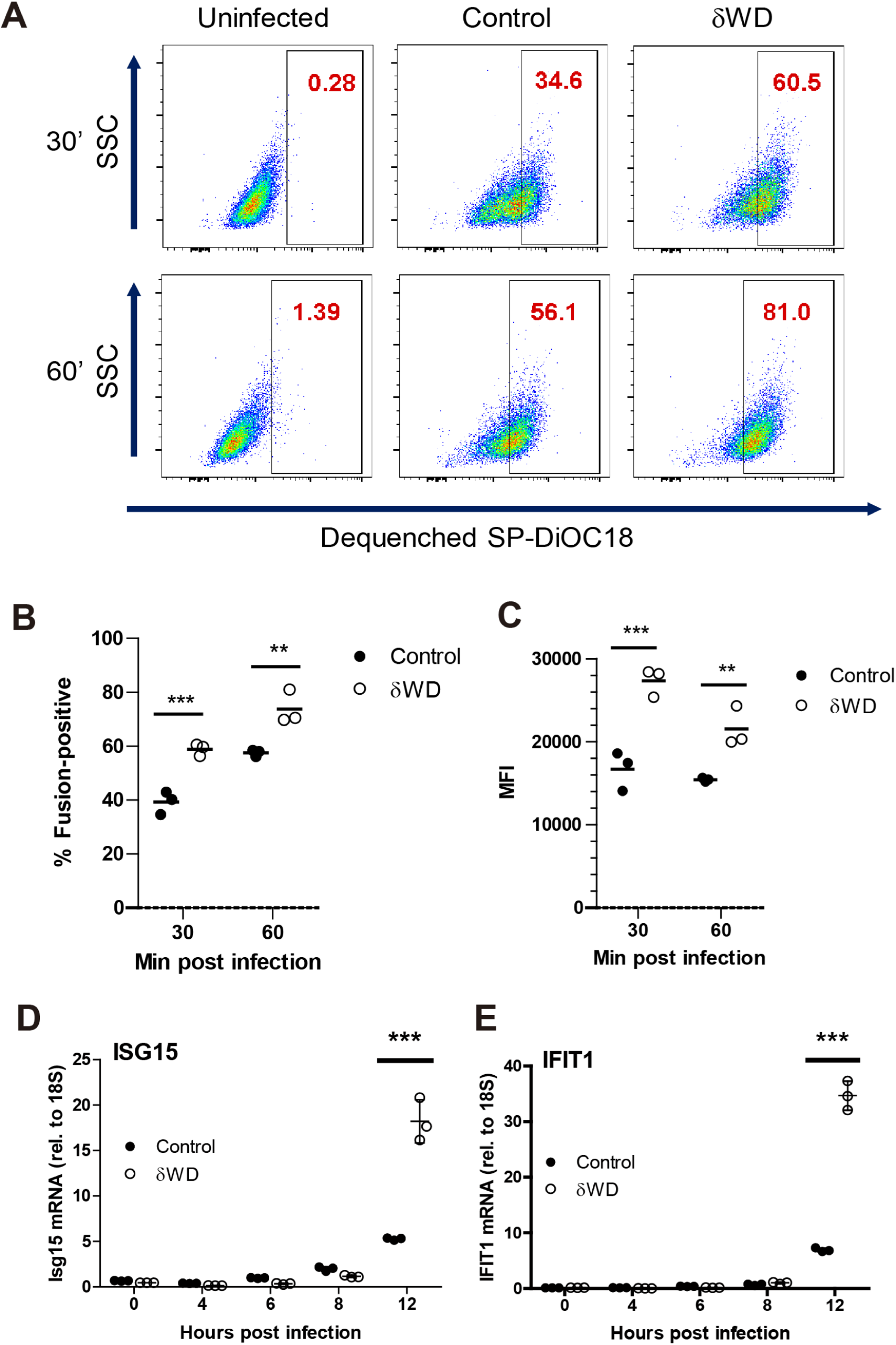
Non-canonical autophagy reduces endosome fusion *in vitro* and reduces interferon signalling. Panels **A-C**, Sub-confluent monolayers of MEFs from δWD or littermate control mice were incubated with dual-labelled (SP-DiOC18/R18) IAV at 40C for 45 min and warmed to 37°C for increasing times. Cells were harvested by trypsinisation, fixed in PFA and analysed by flow cytometry. **(A)**. Representative plots showing de-quenched SP-DiOC18. Numbers in the gates show percentage of cells positive for fusion as determined by comparison to uninfected cells. **(B)** Percentage of cells positive for fusion **(C)** Median fluorescence intensity of dequenched SP-DiOC18 signal. **B** and **C** show individual replicates with a bar at the mean and were compared using two-way ANOVA with Bonferroni post-tests. pH-dependent fusion was assessed by adding bafilomycin A1 to the infection assay (Fig S9). Panels **D** and **E**, MEFs from δWD or littermate control mice were infected with IAV. At the indicated time points, mRNA transcripts were evaluated by qPCR for (**B**) ISG15 and (**D**) IFIT1. Data from 3 replicates are shown, bars represent the mean ± SD and were compared by two-way ANOVA with Bonferroni post-tests.

## Discussion

Respiratory viruses such as influenza A virus (IAV) and SARS CoV-2 can move from animal reservoirs to create human pandemics with high morbidity and mortality. The danger posed by pandemic spread of respiratory viruses underlines an urgent need to understand how the airways defend against viral infection. In this study we have analysed the role played by non-canonical autophagy in defending the respiratory tract against infection by IAV *in vivo*. Mice with systemic loss of non-canonical autophagy (δWD) showed profound sensitivity to infection by a low-pathogenicity murine-adapted IAV (A/X31) leading to extensive viral replication throughout the lungs, dysregulated cytokine production, fulminant pneumonia and lung inflammation leading to high mortality and death usually seen after infection with virulent strains (Belser et al., 2011). These signs mirror the cytokine storms and mortality seen in humans infected with highly pathogenic strains of IAV such as the 1918 ‘Spanish’ Influenza (Belser et al., 2011).

The observation that bone marrow transfers from wild-type mice were unable to protect δWD mice from IAV suggested that protection against IAV infection *in vivo* was independent of leukocytes and did not require non-canonical autophagy in leukocyte populations (e.g. macrophages, dendritic cells, neutrophils, granulocytes, lymphocytes). In a reciprocal experiment the linker and WD domains of ATG16L1 were deleted specifically from myeloid cells. These mice, which lack non-canonical autophagy in phagocytic cells (LAP), but maintain non-canonical autophagy in other tissues, failed to show increased sensitivity to IAV infection. Thus, protection against severe IAV-associated disease in the respiratory tract of the host relies heavily on non-canonical autophagy in non-leukocyte populations.

Activation of non-canonical autophagy in phagocytic cells leads to LC3 associated phagocytosis (LAP) where TLR signalling and reactive oxygen species (ROS) recruit LC3 to phagosomes. A lack of involvement of LAP in protection against IAV disease *in vivo* was surprising because the activation of LAP in phagocytic cells such as macrophages, dendritic cells and neutrophils by would provide a powerful means of recognising and controlling microbial infection *in vivo. In vitro* studies show that activation of acid sphingomyelinase by *Listeria monocytogenes* (Gluschko et al., 2018) and subsequent ROS production by NOX2 recruit LC3 to phagosomes. Similarly, activation of TLR2 and NOX2 by *Legionella dumoffii in vitro* signal ULK1-independent translocation of LC3 to single-membraned vacuoles containing Legionella (Hubber et al., 2017). In both cases LC3 promotes fusion with lysosomes. The observation that virulence factors such as the GP63 metalloprotease of *Leishmania major* and melanin of *Aspergillus fumigatus* prevent recruitment of NOX2 to phagosomes to prevent LAP (Akoumianaki et al., 2016, Kyrmizi et al., 2018, Matte et al., 2016) further suggest that non-canonical autophagy in phagocytes should provide a defence against infection. One reason for the discrepancy may be that the studies cited above have focused on *in vitro* experiments using microbes with a tropism for macrophages, rather than *in vivo* studies where pathogens encounter epithelial barriers.

Intranasal infection of mice with IAV results in rapid infection of principally airway and pulmonary epithelial cells (Akram et al., 2018). The results of *in vivo* challenge of radiation chimaeras and δWD^phag^ mice strongly suggest that non-canonical autophagy in the epithelium rather than leucocytes is responsible for restricting IAV infection. This was supported by *ex vivo* experiments where virus titres and interferon responses were 5 - 10 fold greater in precision-cut lung slices, and MEFs from δWD mice. Furthermore, loss of non-canonical autophagy increased fusion of IAV envelope with endosomes, and increased activation of interferon signalling pathways. Both assays suggest that non-canonical autophagy reduces IAV entry and delivery of viral RNA to the cytoplasm. This would explain reduced interferon signalling, and at the same time the delayed escape of IAV into the cytoplasm would increase the transfer of endocytosed virus to lysosomes for degradation. The precise mechanisms employed by non-canonical autophagy to reduce virus entry from endosomes are unknown. This may involve recruitment of LC3 to endosomes by TMEM59 (Boada-Romero et al., 2016) to increase fusion with lysosomes, or by maintaining membrane repair during virus entry, as observed for bacteria such as *S*. Typhimurium and *Listeria monocytogenes* (Kreibich et al., 2015, Tan et al., 2018). A p22^phox^-NOX2 pathway that recruits LC3 to vacuoles containing *S*. Typhimuriumin epithelial cells (Huang et al., 2009) may also be activated during IAV entry and hamper lethal infection.

δWD mice infected with IAV appeared to be unable to resolve inflammatory responses resulting in sustained expression of pro-inflammatory cytokines, morbidity and a striking lung pathology characterized by profuse migration of neutrophils into the airway at day 3 followed by macrophages on day 7. pDCs detect IAV-infected cells and produce large amounts of cytokines, in particular IFNαβ, that in severe infections can enhance disease(Davidson et al., 2014). The fact that morbidity in δWD mice could be decreased by depleting pDCs indicates that excessive cytokine production, amplified by pDCs was a major factor. This is not due to a lack of non-canonical autophagy/LAP in pDC as bone-marrow chimaeras of δWD mice with wild-type leukocytes have the same phenotype as δWD mice. IAV is recognized by endosomal TLR3 in respiratory epithelial cells and RIG-I detects virus replicating in the cytosol leading to activation of IRF3 and NFkB with subsequent induction of interferon, ISG and proinflammatory cytokine production (Iwasaki & Pillai, 2014). Increased inflammation may result directly from increased virus in the lungs, but the increased fusion of IAV envelope with endosomes in δWD mice may increase delivery of viral RNA to the cytoplasm resulting in the sustained pro-inflammatory cytokine signalling. A similar pro-inflammatory phenotype resulting from decreased trafficking of inflammatory cargoes is observed following disruption of non-canonical autophagy by LysMcre-mediated loss of Rubicon from macrophages or microglia (Heckmann et al., 2019, Martinez et al., 2016).

We have dissected the roles played by conventional autophagy and non-canonical autophagy *in vivo* by removing the linker and WD domain from ATG16L1 to prevent conjugation of LC3 to single-membraned endo-lysosome compartments (Rai et al., 2019). An alternative approach has been to target pathways upstream of LC3 conjugation where deletion of Rubicon produces a selective block in LAP (Heckmann et al., 2019, Martinez et al., 2015). Rubicon stabilises the PHOX:NOX2 complex (Yang et al., 2009) allowing reactive oxygen species (ROS) to induce binding of ATG16L1 to endo-lysosome membranes (Martinez et al., 2015). Mouse models relying on loss of Rubicon show defects in the clearance of bacterial and fungal pathogens and apoptotic cells (Martinez et al., 2016, Martinez et al., 2015), but have not yet been studied in the context of viral infection. Furthermore, disruption of Rubicon leads to upregulation of IL-1β, IL6 and TNF-α secretion, and the mice fail to gain weight and develop an autoimmune disease that resembles Systemic Lupus Erythematosus (Heckmann et al., 2017, Martinez et al., 2016). This exaggerated inflammation might make it difficult to predict if any altered responses to infection observed in Rubicon-/-mice, particularly lung inflammation, resulted directly from loss of non-canonical autophagy, or from upstream changes in cytokine regulation caused by loss of Rubicon.

Several non-canonical pathways leading to recruitment of LC3 to endo-lysosomal compartments, rather than phagosomes, are beginning to emerge. Non-canonical autophagy in microglia facilitates endocytosis of β−amyloid and TLR receptors to reduce β-amyloid deposition and inflammation in mouse models of Alzheimer’s disease (Heckmann et al., 2019). This may involve and interaction between the WD domain and TMEM59 which is required for β-amyloid glycosylation (Ullrich et al., 2010). Lysosomotropic drugs, which stimulate direct recruitment of LC3 to endosomes, create pH and osmotic changes that may mimic the consequences of viral infections that perturb endosome membranes or deliver viroporins to endo-lysosome compartments. It will be interesting to see if the WD and linker domains of ATG16L1 limit infection by other microbes at epithelial barriers *in vivo*, particularly infection of the respiratory tract by SARS-CoV-2. This may be true for picornaviruses where LC3 is recruited to enlarged endosomes during entry of Foot and Mouth Disease virus (Berryman et al., 2012) and following LC3 accumulation on megaphagosomes in pancreatic acinar cells during coxsackievirus B3 infection (Kemball et al., 2010). In the specific cases of IAV and SARS-Cov-2, non-canonical autophagy at epithelial barriers is likely important for innate control of new pathogenic strains, where acquired immunity from previous infection may be absent or less effective. It will be valuable to assess whether human allelic variants of ATG16L1 confer altered resistance/susceptibility to respiratory infections such as IAV and whether drug-based manipulation of non-canonical autophagy can increase resistance at the respiratory epithelial barrier.

## Acknowledgements

This work was funded in part by Biotechnology and Biological Sciences Research Council (BBSRC) grant BB/R00904X/1 to JPS, JLC SRC, PPP, UM and TW, and through BBSRC Institute Strategic Programme Gut Microbes and Health BB/R012490/1: BBS/E/F/000PR10353, BBS/E/F/000PR10355

## Author contributions

TW, JPS, PPP, SRC and JLC conceived the experiments. Mouse strains were generated by UM and genotyped by MJ, SR and WZ. Immunological homeostasis was assessed by WZ, AM and AZ. Animal infections were carried out by JPS, YW, WZ, and PS and histology and immunohistology by AK. IAV fusion was analysed by YY with BB. Downstream analysis was performed by YW, WZ, PS, TP and PPP. *In-vitro* analysis was performed by BB, TP, YY and PPP. The manuscript was drafted by TW, JPS, YB, PPP, RAT and UM and edited and approved by all authors

## Declaration of interests

Authors declare no competing interests.

## Materials and Methods

### Cell culture and virus

Influenza virus A/HKx31 (X31, H3N2) was propagated in the allantoic cavity of 9-day-old embryonated chicken eggs at 35° C for 72 h. Titres were determined by plaque assay using MDCK cells with an Avicel overlay.

### Mice

All experiments were performed in accordance with UK Home Office guidelines and under the UK Animals (Scientific procedures) Act1986.

The generation of δWD mice (*Atg16L1*^δWD/δWD^) has been described previously (Rai et al., 2019). Generation of δWD^phag^ and *Atg16L1*^fl/fl^-LysMCre mice is described in detail in Fig. S7. Comparisons were made using age and sex-matched littermate control mice for each individual genotype. Generation and breeding of mice was approved by the University of East Anglia Animal Welfare and Ethical Review Body and performed under UK Home Office Project License 70/8232.

Influenza Infection studies were performed at the University of Liverpool, approved by the University of Liverpool Animal Welfare and Ethical Review Body and performed under UK Home Office Project License 70/8599. Studies used 2-3 m old male and female mice that had been back-crossed to C57BL/6J. Mice were maintained under specific pathogen-free barrier conditions in individually ventilated cages (Greenline GM500, Techniplast) at a temperature of 22°C (± 2°C), humidity 55% (± 10%), light/dark cycle 12/12 hours (7 am to 7 pm), food CRM(P) and RO or filtered water *ad lib*. Colonies were screened using the Charles River surveillance plus PRIA health screening profile every 3 months to ensure SPF status.

For IAV infection, animals were randomly assigned into multiple cohorts, anaesthetised lightly by the i.m. route with 150 mg/kg ketamine (Ketavet, Zoetis UK Ltd) and separate cohorts inoculated intra-nasally with 10^3^ PFU IAV strain X31 in 50 µl sterile PBS. Mice were infected between 9 and 11 AM. Animals were sacrificed at variable time-points after infection by cervical dislocation. Tissues were removed immediately for downstream processing. Sample sizes of n = 6 were used as determined using power calculations and previous experience of experimental infection with these viruses. For survival analysis, a humane endpoint was determined using a scoring matrix that included excessive (>20%) weight loss.

To specifically deplete plasmacytoid dendritic cells (pDCs), mice were treated with anti-PDCA-1 (Cambridge Bioscience) or IgG2b isotype-matched control, using a dose of 500 mg per 200 ml via the i.p. route on day 1 of infection with IAV and every 48 h thereafter (Davidson et al., 2014).

### Generation and analysis of radiation chimeras

The general strategy is shown in Fig S6A. Mice were subjected to whole body irradiation with 11 Gy in two doses 4 h apart using a ^137^Cs source in a rotating closed chamber. Bone marrow was collected from male wild-type C57BL/6-Ly5.1 (B6.SJL-*Ptprc*^*a*^*Pepc*^*b*^/BoyCrl; Atg16L1^+/+^) mice that are congenic for the CD45.1 allele or from δWD mice (that are congenic for CD45.2). The C57BL/6 CD45.1 marrows were used to enable confirmation of chimaerism by FACS analysis of bon-marrow-derived cells as littermate control and δWD mice are CD45.2 (Fig S6B). The femur and tibia of the donor mouse was collected and sterilised for 2 min in 70% ethanol. The ends of the bones were removed and PBS was used to flush out the bone marrow through a 40 μm cell sieve. Red blood cell lysis was performed using 0.83% ammonium chloride and the cells were washed twice in PBS and re-suspended at a concentration of 10^7^ cells/ml. T cell depletion was performed prior to transfusion by using a commercial mouse hematopoietic progenitor cell isolation kit (EasySep, STEMCELL™ Technologies, #19856).

After depletion, 10^6^ donor bone marrow cells were injected into each irradiated mouse by tail vein injection 3 h following irradiation. Mice were then allowed to recover for 12 weeks with daily monitoring of mouse weights and general condition for at least the first two weeks to monitor for any severe radiation sickness or illness due to being immunocompromised.

For chimaerism analysis, approximately10^6^ spleen cells were analysed by flow cytometry using fluorochrome-conjugated monoclonal antibodies specific for CD45.1 (clone A20 eBioscience), CD45.2, (clone 104 eBioscience). As shown in Fig. S6B, in the groups where CD45.1 marrow was transplanted all mice were >95% chimaeric.

### Flow Cytometric analysis of Cells

Brocho-alvolear lavage fluid (BAL) was obtained by lavage of mice via the trachea using 1 ml ice-cold RPMI containing 5% FCS. For lung tissue, single-cell suspensions were made from minced lung and subjected to collagenase and DNase I digestion, then treated with ACK buffer to remove red blood cells. In both cases, approximately10^6^ cells were incubated in 100 µl of Fc block (clone 2.4G2, BD Bioscience) diluted in PBS, 2% FCS (PBS-FCS) for 15 min at 4 **°**C prior to the addition of fluorochrome-conjugated monoclonal antibodies and incubation for 30 min at 4**°**C in the dark. Cells were then washed in PBS-FCS, fixed in 4% paraformaldehyde in PBS for 15 min at 20°C prior to analysis on a MACSQuant Analyzer 10 (Miltenyi Biotech UK). Data were analysed using FlowJo (FlowJo, LLC). Antibodies used included: CD45, Ly6G, CD11c, CD11b, F4/80 (all eBioscience). Neutrophil populations in BAL were identified as CD45^+^, CD11c^-^, CD11b^+^, Ly6G^+^. Macrophage/monocyte populations in lung tissue were identified as CD45^+^, CD11c^-^, CD11b^+^, F4/80^+^.

### Histology, immunohistochemistry

Tissues were fixed in 4% buffered paraformaldehyde (PFA; pH7.4) for 24 h and routinely paraffin wax embedded. Consecutive sections (3-5 µm) were either stained with haematoxylin and eosin (HE) or used for immunohistochemistry (IH).

IH was performed to detect influenza antigens and to identify neutrophils and neutrophil extracellular traps (NETs) and macrophages using the horseradish peroxidase (HRP) and the avidin biotin complex (ABC) method. The following primary antibodies were applied: goat anti-IAV (Meridian Life Sciences Inc., B65141G), rat anti-mouse Ly6G (clone 1A8, Biolegend; neutrophil marker), rabbit anti-Iba-1 (antigen: AIF1; Wako Chemicals; microglia/macrophage specific marker), and rabbit anti-histone H3 (citrulline R2 + R8 + R17; Abcam; NET marker). Briefly, after deparaffination, sections underwent antigen retrieval in citrate buffer (pH 6.0, 20 min at 98°C) followed by blocking of endogenous peroxidase (peroxidase block, S2023, Dako) for 10 min at room temperature (RT). Slides were then incubated with the primary antibodies (diluted in dilution buffer, Dako) for a) Iba-1 (60 min at RT), followed by a 30 min incubation at room temperature with the secondary antibody (Envision mouse and rabbit, respectively, Dako) in an autostainer (Dako), and b) Ly6G (60 min at RT), followed by rabbit anti-rat IgG and the ABC kit (both 30 min at RT; Ventana). Staining for histone H3 was undertaken with an autostainer (Discovery XT, Ventana), using citrate buffer, dilution buffer and detection kits provided by the manufacturer. The antibody reaction was visualized with 3,3’-diaminobenzidine and sections counterstained with haematoxylin.

### Statistical analysis

Data were analysed using the Prism package (version 5.04 Graphpad Software). *P* values were set at 95% confidence interval. A repeated-measures two-way ANOVA (Bonferroni post-test) was used for time-courses of weight loss; two-way ANOVA (Bonferroni post-test) was used for other time-courses; log-rank (Mantel-Cox) test was used for survival curves; one-way ANOVA (Tukey’s post-hoc) was used to compare three or more groups side-by-side; Mann-Whitney *U* test was used to compare two groups. All differences not specifically stated to be significant were not significant (p > 0.05). For all figures, *p < 0.05, **p <0.01, ***p < 0.001, ****p < 0.0001.

### Cells and cell culture

Mouse embryonic fibroblasts (MEFs) were generated by serial passage of cells taken from mice at embryonic day 13.5 and cultured in DMEM (ThermoFisher scientific, 11570586) with 10% FCS. Bone marrow derived macrophages (BMDMs) were generated from femur and tibia flushed with RPMI-1640 (Sigma, R8758). Macrophages were generated from adherent cells in RPMI-1640 containing 10% FCS and M-CSF (Peprotech, 315-02) (30 ng/ml) for 6 d. Macrophage populations were quantified by FACS using antibodies against CD16/CD32, F4/80 and CD11b (BioLegend, 101320, 123107).

### Precision-cut lung slices

Infection of *ex vivo* lung slices was used to examine the responses of lungs without any contribution from recruited leukocytes, which could not be present. Mouse lungs were inflated with 2% low melting point agarose in HBSS and then sliced into 300 µm sections using a vibrating microtome. They were then cultured overnight in DMEM/F12 medium (Thermofisher 21331020) prior to infection with IAV.

### IAV Endosome fusion assay

The envelope of purified IAV (0.1mg protein mL^-1^) was labelled using an ethanol solution containing 33µM 3,3’-dioctadecyloxacarbocyanine (DIOC18) and 67µM octadecyl rhodamine B (R18). Aggregated virus was removed by a 0.22µm filter (Millipore). Sub-confluent monolayer of MEFs cultured in DMEM, (50mM HEPES, 0.2% BSA) were incubated with the labelled virus at 4°C for 45 min and warmed to 37°C for increasing times. Cells were harvested by trypsinisation and fixed in 4% PFA for 20 min. Cell pellets (2500rpm, 4 min) were resuspended in 100µL FACS buffer (1xPBS 1%BSA) and analysed by FACs using a NovoCyte Flow cytometer FlowJo software. pH independent fusion was assessed by adding bafilomycin A1 to the infection assay.

### Autophagy and non-canonical autophagy

Autophagy was activated by incubating cells in Hanks balanced salt solution (HBSS) (ThermoFisher, 11550456) for 2 h at 37°C. Non-canonical autophagy was stimulated in BMDMs by incubation with Zymosan A (Alexa Fluor 594-labelled; Thermofisher Z23374). LC3 was analyzed by immunofluorescence microscopy and LC3i:LC3II Western blot.

### qPCR for cytokine transcription

Lung lobes were snap frozen and homogenized using a TissueLyser (Qiagen). Tissue culture cells were washed twice using PBS. Total RNA was extracted by Trizol-chloroform (Thermofisher 15596018) and purified by RNeasy MinElute cleanup kit (Qiagen 74204). RNA was analysed by qPCR using SYBR Green/7500 (Thermofisher S7563) standard Real-Time PCR system (Applied Biosystems, Grand Island, NY) and primer sets as detailed in Table S1. Relative amounts of mRNA expression were normalized to 18S rRNA.

**Table 1.** Primer sequences for mRNAs analysed by RT-qPCR.

| Target | Catalogue number<sup>†</sup> |
| --- | --- |
| ISG15 | QT00322749 |
| IFIT1 | QT01161286 |
| IL-1β | QT01048355 |
| TNF-α | QT00104006 |
| CXCL1 | QT00115647 |
| CCL2/MCP-1 | QT00167832 |
| 18S ribosomal RNA | QT02448075 |

<sup>†</sup> Catalogue numbers refer to validated QuantiTect primer sets (Qiagen)

### Western blotting

Cells were lysed using M-PER reagent (ThermoFisher 78501) with complete protease inhibitor cocktail (Sigma, 04693159001) and clarified by centrifugation. Extracted proteins (20 µg) were separated on a precast 4–12% gradient SDS-PAGE gels (Expedeon, NBT41212), transferred to immobilon PVDF (Millipore, IPFL00010) and probed using antibodies for ATG16L1 (MBL M150-3), LC3A/B (Cell signaling 41085) and actin (Sigma, A5441). Primary antibodies were detected using IRDye labelled secondary antibodies (LI-COR biosciences, 926-32211, 926-68020) and visualised by Odyssey infrared system (LI-COR).

### Fluorescence imaging

Cells were fixed in ice cold methanol and non-specific binding was blocked using 5% goat serum plus 0.3% Triton-X100 in PBS followed by incubating with anti LC3A/B (Cell Signalling 4108) or anti-ATG16L1 (MBL M150-3). Cells were washed and then incubated with anti-rabbit-Alexa 488 (Thermofisher 10729174). After washing, cells were counterstained with 4’, 6 diamidino-2-phenylindole (DAPI) (ThermoFisher scientific, 10116287) and mounted with Fluoromount-G (Cambridge Bioscience). Cells were imaged on a Zeiss Imager M2 Apotome microscope with a 63x, 1.4 NA oil-immersion objective.

## Supplementary Materials for

**Fig. S1.**
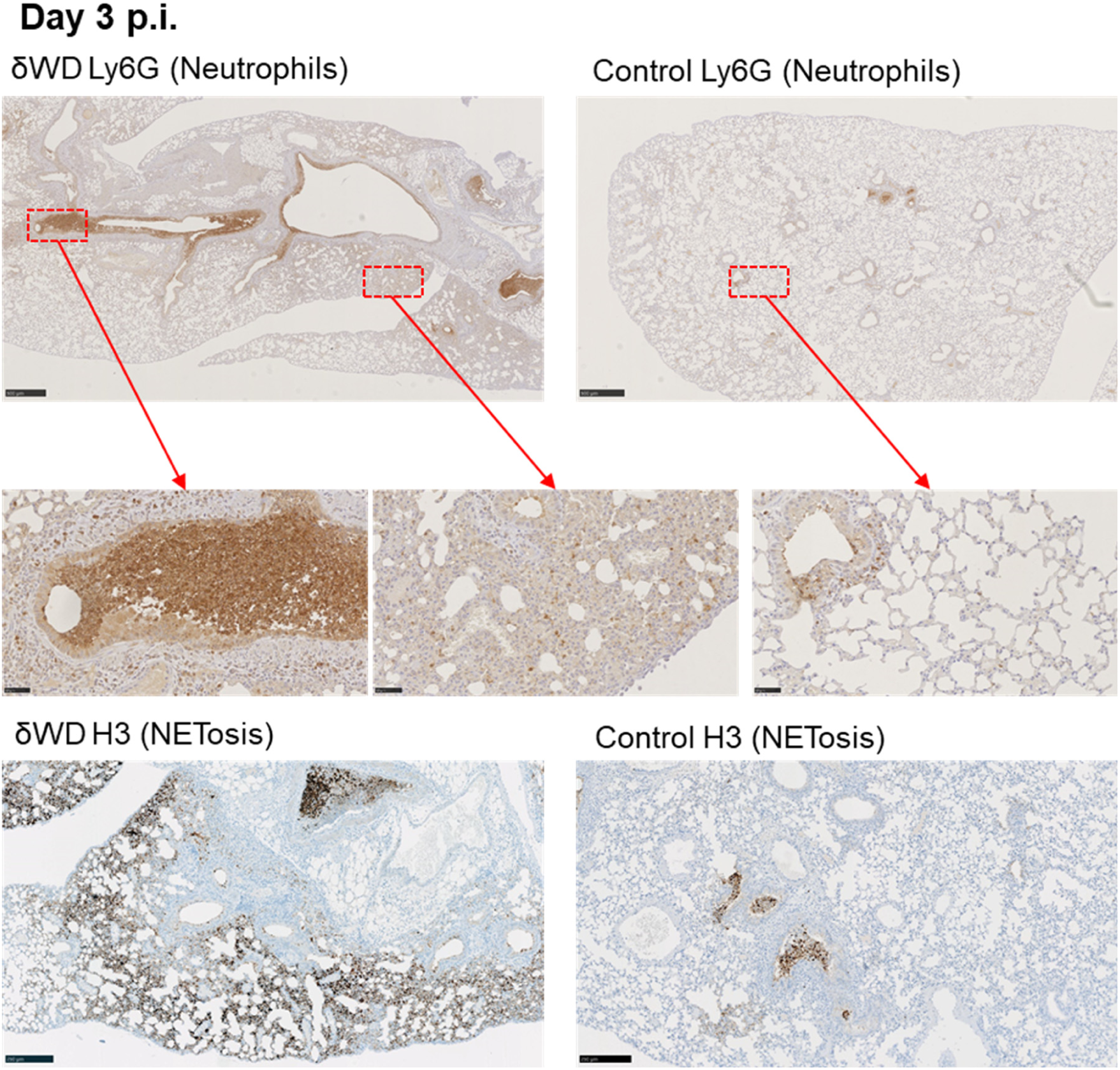
Increased neutrophilia and NETosis in IAV-infected mice deficient in non-canonical autophagy. δWD and littermate control mice were infected i.n. with 10^3^ pfu IAV X31. Lung tissues were harvested at 3d p.i. Neutrophils and H3 (marker of NETosis) were detected by IH using anti-Ly6G and anti-H3, visualized with DAB and counter-stained with hematoxylin. Micrographs of representative areas from lungs of six mice are shown. Scale bars represent 500 µm (upper panels), 50 µm (middle panels) or 250 µm (lower panels). There are dramatically increased numbers of neutrophils in airways (bronchi and bronchioles) and lung parenchyma of δWD mice, accompanied by markedly-increased NETosis, indicating significant neutrophil degeneration.

**Fig. S2.**
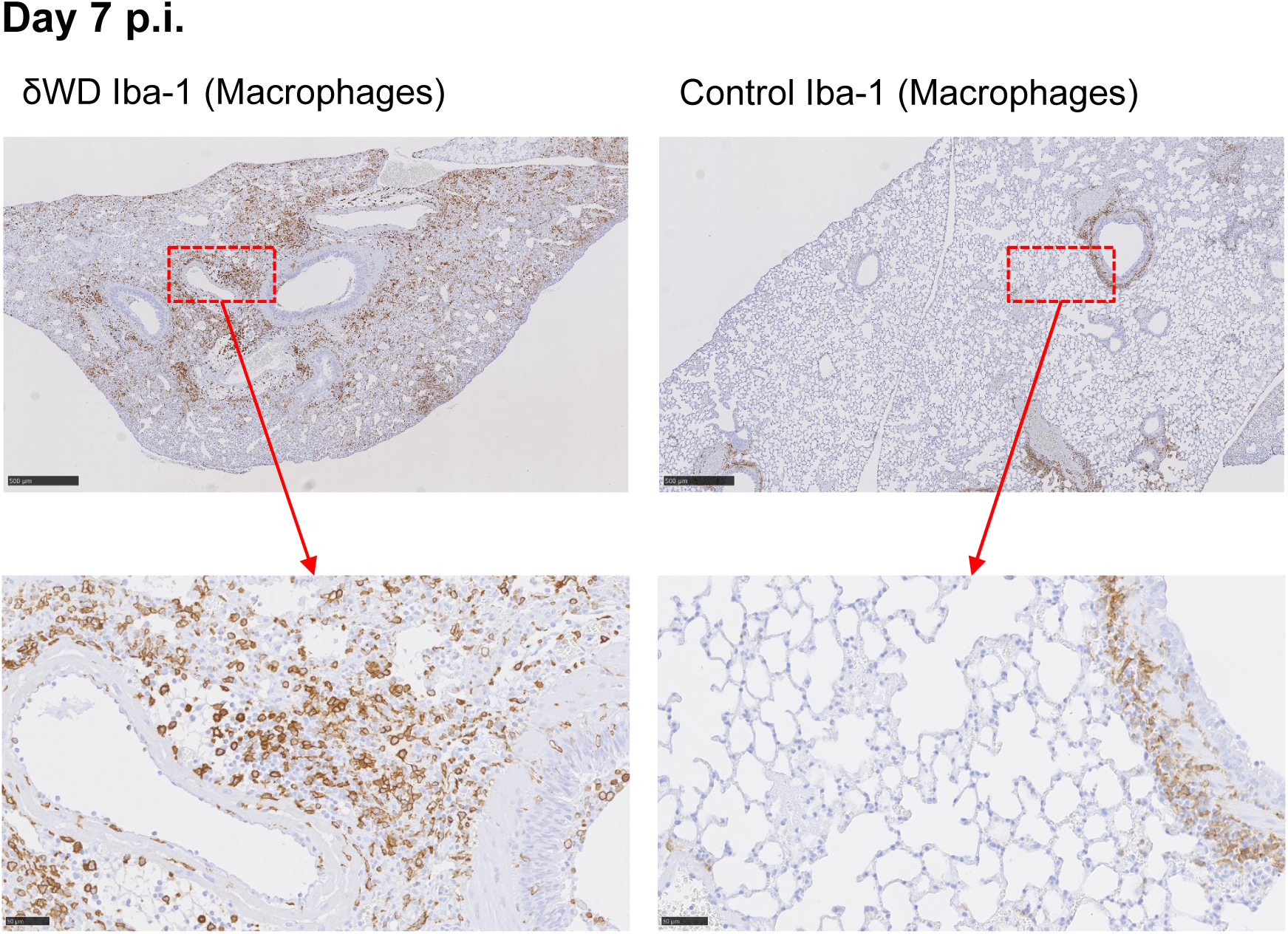
Increased macrophage rich inflammation in IAV-infected mice deficient in non-canonical autophagy. δWD and littermate control mice were infected i.n. with 10^3^ pfu IAV X31. Lung tissues were harvested at 7d p.i. Macrophages were detected by IH using anti-Iba-1, visualized with DAB and counter-stained with hematoxylin. Micrographs of representative areas from lungs of six mice are shown. Scale bars represent 500 µm (upper panels) and 50 µm (lower panels). Lower panels are the same as in Fig. 2B. Upper panels show the lower magnification images of the lung to illustrate the general nature of the observations. There is clearly increased inflammation in δWD mice with higher numbers of macrophages in the lung parenchyma.

**Fig. S3.**
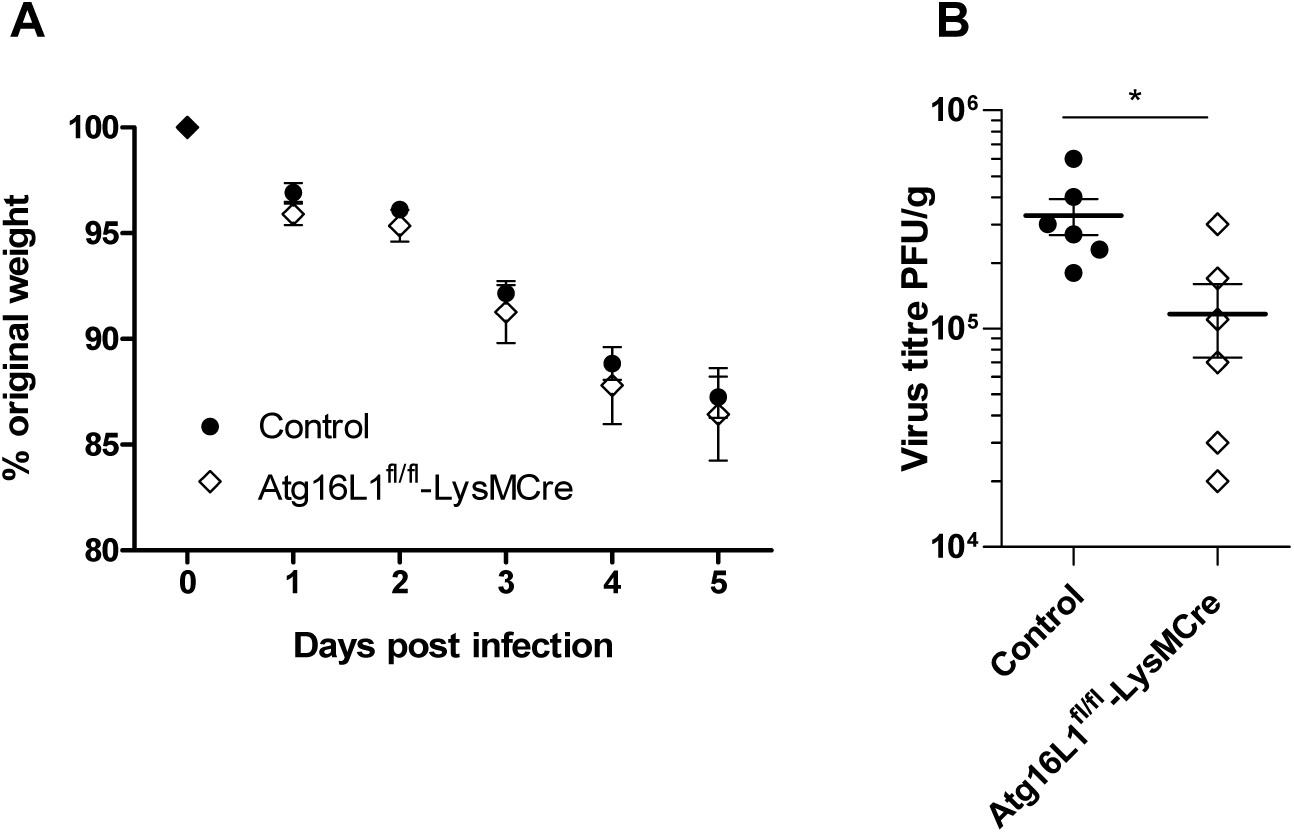
LysMcre-mediated loss of canonical macro-autophagy from phagocytes decreases sensitivity to IAV infection. *Atg*16L1^fl/fl^-LysMcre mice and littermate controls; *n* = 5 or 6 per group) were infected i.n. with 10^3^ pfu IAV X31. **Panel A**. Mice were weighed daily and the weights presented as a percentage of the starting weight. **Panel B**. Lung tissues were taken at 5 d.p.i. and virus titer determined by plaque assay. Data represent the mean value ± SEM. Analysis using the Mann-Whitney U test showed a significant difference (* p < 0.05). Thus, *Atg*16L1^fl/fl^-LysMcre mice that are deficient in canonical autophagy in phagocytes lose weight at the same rate as littermate controls but are more resistant to virus replication as they have lower lung virus titres.

**Fig. S4.**
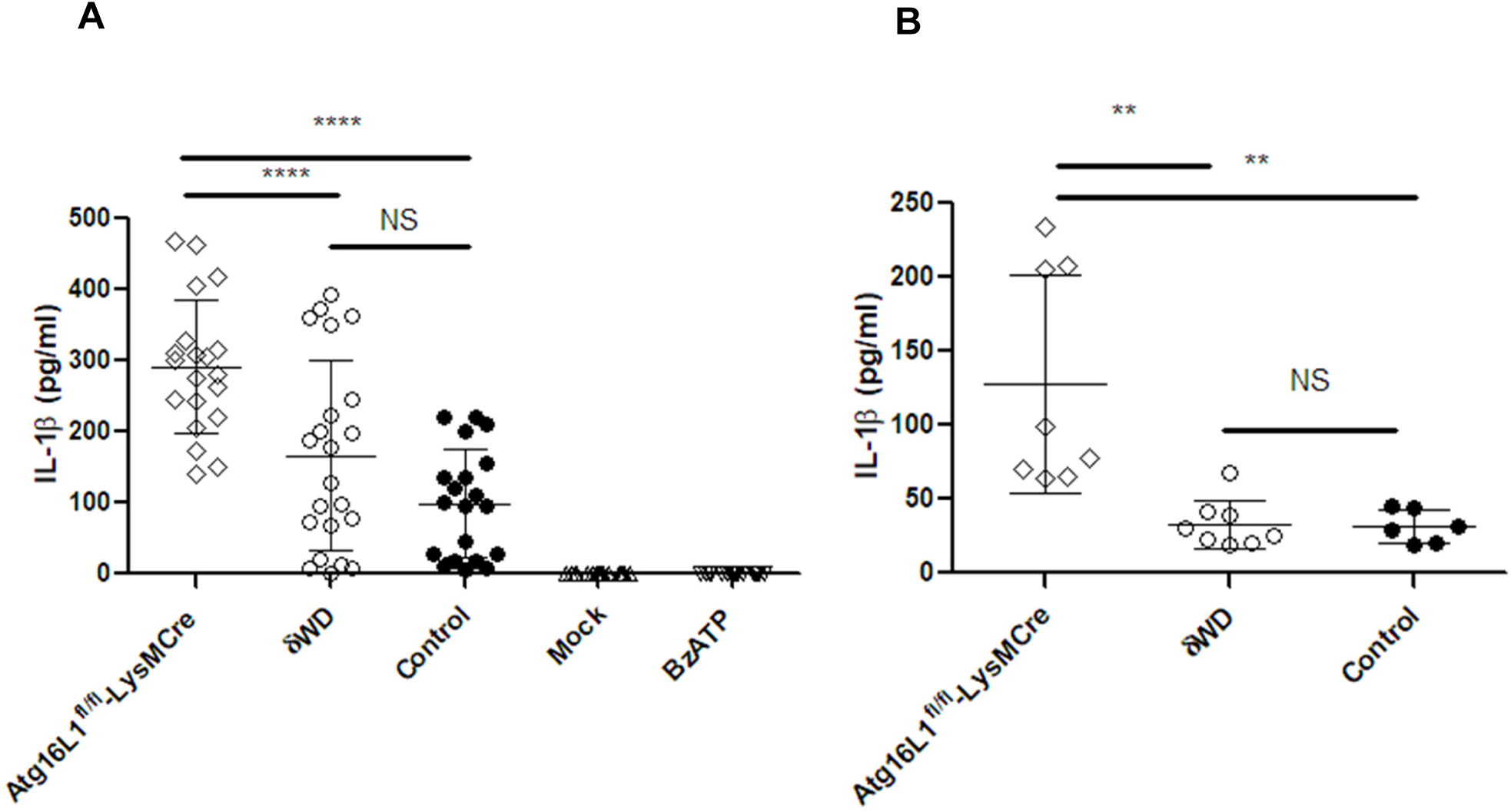
Mice deficient in non-canonical autophagy do not have elevated IL-1β in response to LPS stimulation. LysMcre-mediated deletion of autophagy genes from mice leads to increased inflammatory threshold characterised by raised secretion of IL-1β from macrophages (*15*), and in the lung this can increase resistance to IAV infection (*16*). The possibility that the δWD mutation could affect IL-1β secretion was tested by stimulating BMDM with bacterial lipopolysaccharide (LPS) and BzATP (P2X_7_ receptor agonist) or challenging mice with LPS. **Panel A**. Bone marrow-derived macrophages (BMDM) from mice strains as indicated were incubated with 100 ng/ml of LPS for 4 h and 150 μM of BzATP for 30 min. Supernatants were assayed for IL-1β by ELISA. (Mock group: untreated, BzATP controls only received BzATP). Representative data are shown as the means ± SD of readings from 20 wells per group and were analyzed using one-way ANOVA with Tukey’s post-hoc analysis (**** *p*<0.0001). Approximately three-fold increases in IL-1β secretion were seen for BMDM from Atg16L1^fl/fl^-LysMCre mice. However, IL-1β secretion from δWD BMDM did not differ significantly from littermate controls. **Panel B**. Mouse strains (as indicated) were injected with 20 mg/kg of LPS via the IP route. Serum collected 90 min post injection was assayed for IL-1β by ELISA. In non-treated mice IL-1β was below the detection limit in all 3 strains (not shown). Data are shown as the means ± SD of duplicate assays from 4 mice per group and were analyzed using one-way ANOVA with Tukey’s post-hoc analysis (** *p <* 0.01). Approximately three-fold increases in IL-1β secretion were seen for *Atg*16L1^fl/fl^-LysMCre, however IL-1β secretion for δWD mice did not differ significantly from littermate controls.

**Fig. S5.**
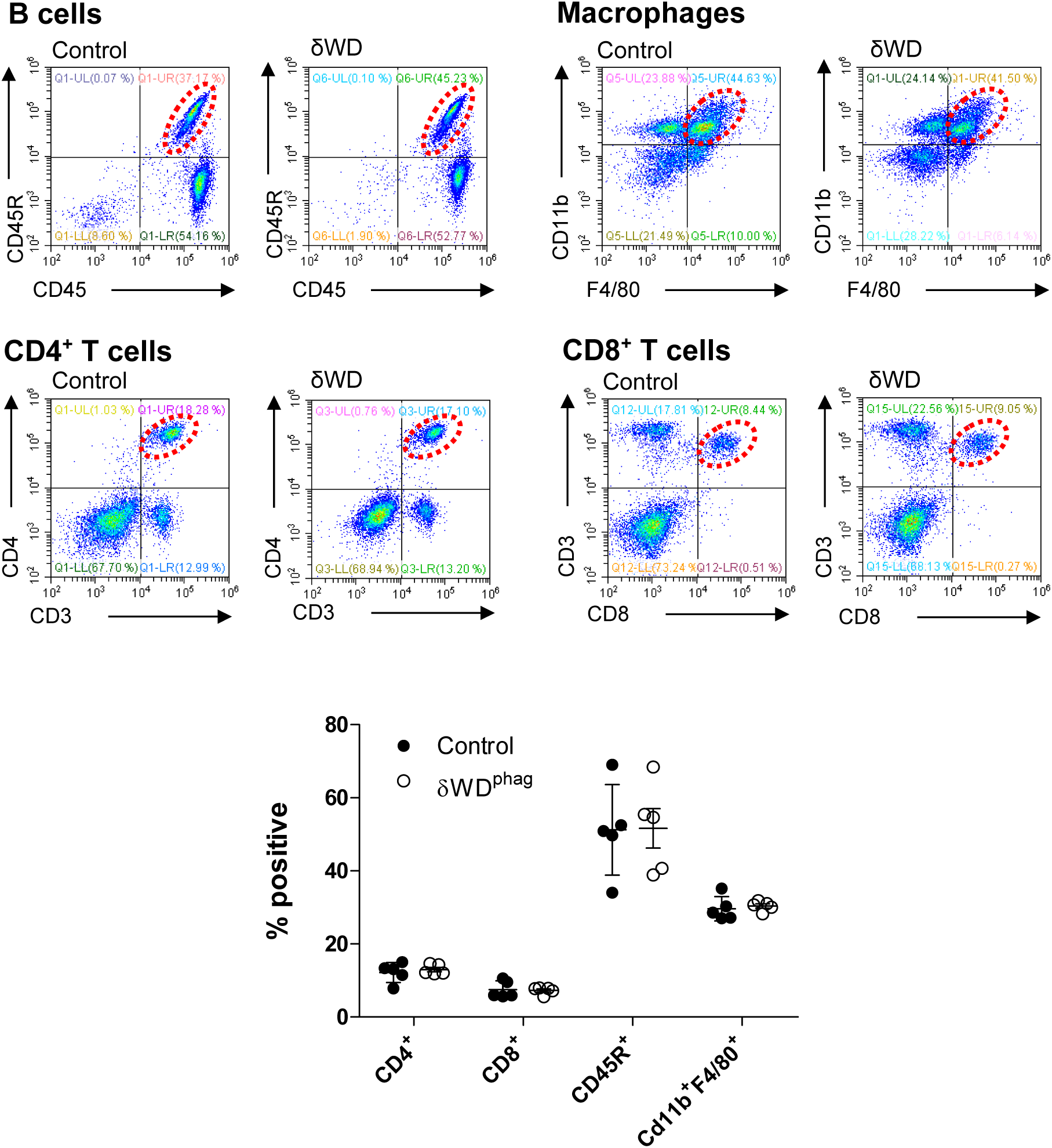
Mice deficient in non-canonical autophagy have normal leukocyte populations. The possibility that the loss of non-canonical autophagy resulted in changes in leukocyte populations was tested by analysing dissociated spleens by FACS using antibodies to T-cell subsets (CD3^+^, CD4^+^ and CD3^+^, CD8^+^), B-cells (CD45R/B220) and macrophages (CD11b, F40/80) Upper panel shows representative FACS profiles from n = 3 mice. Lower panel shows the percentage positive for each population.

**Fig. S6.**
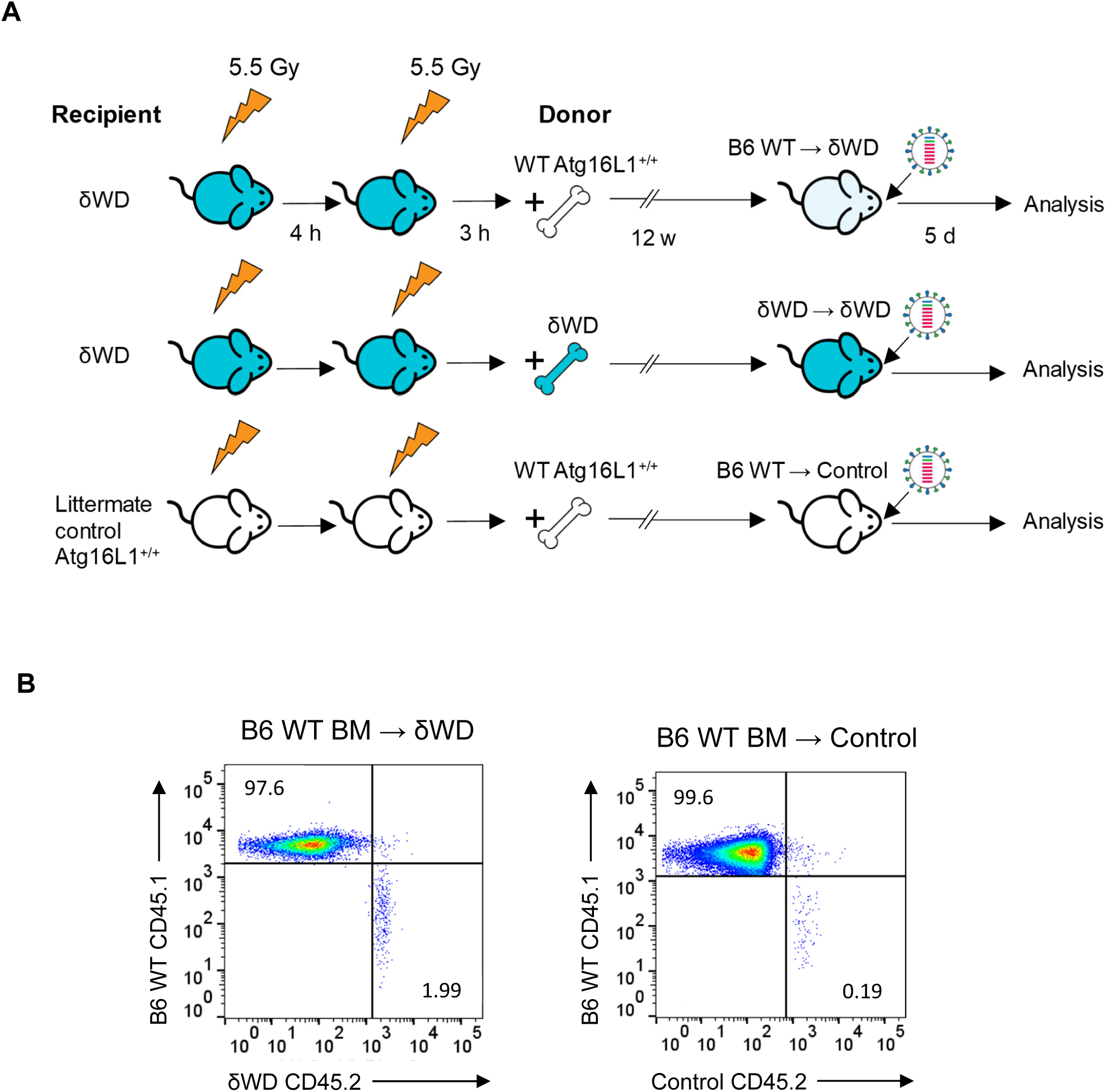
Confirmation of bone marrow transplant radiation chimaerism. **Panel A**. Strategy for making bone-marrow chimaeras. **Panel B**. Chimaerism was confirmed 12 weeks post-transplant in spleen cells by flow-cytometric analysis of congenic markers on leukocytes (CD45.1, CD45.2). Flow plot shows representative plot from one C57BL/6 WT (CD45.1) bone-marrow → δWD (CD45.2) recipient chimaera and one C57BL/6 WT (CD45.1) bone-marrow → littermate control (CD45.2) recipient chimaera. All animals were > 95% chimaeric.

**Fig. S7.**
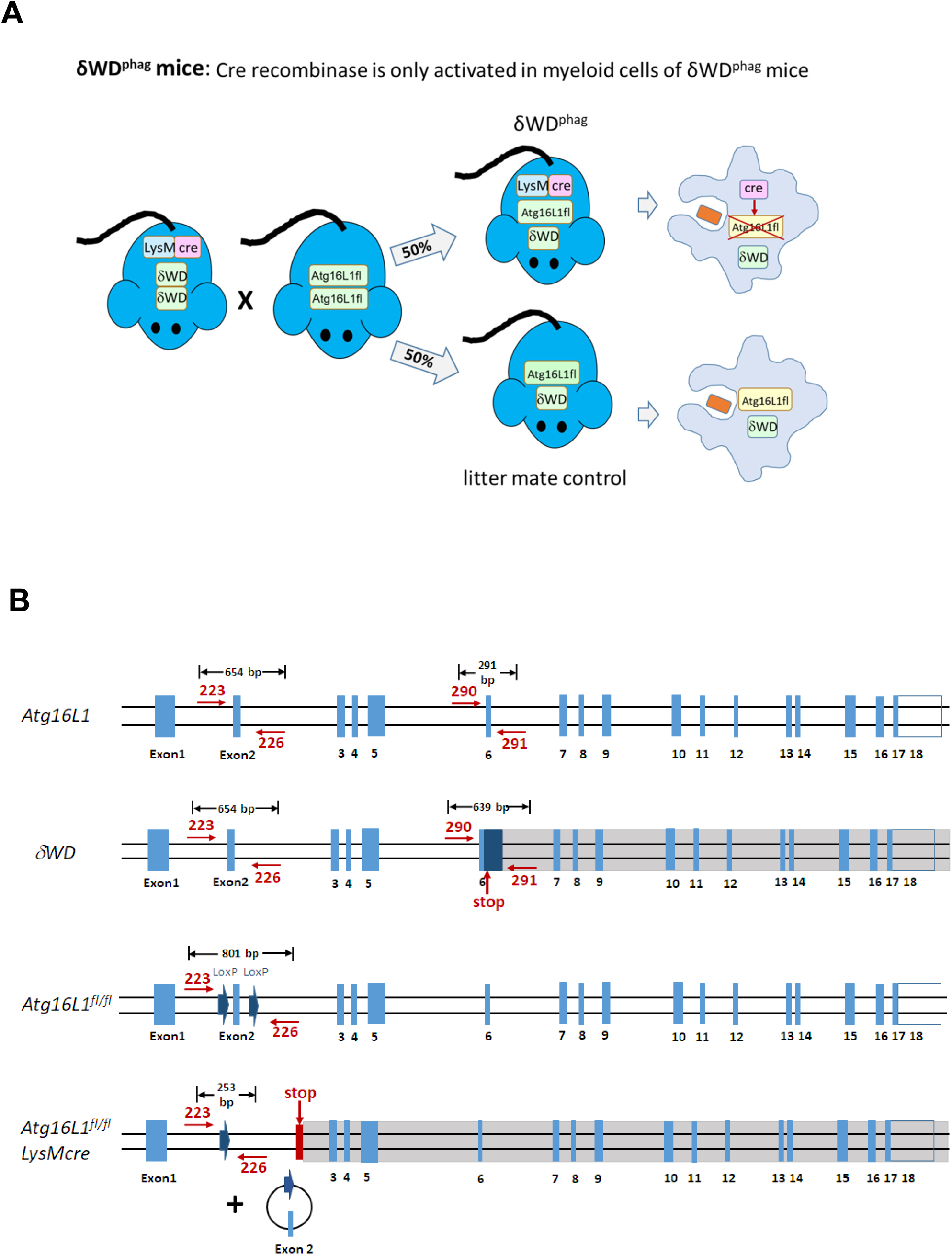
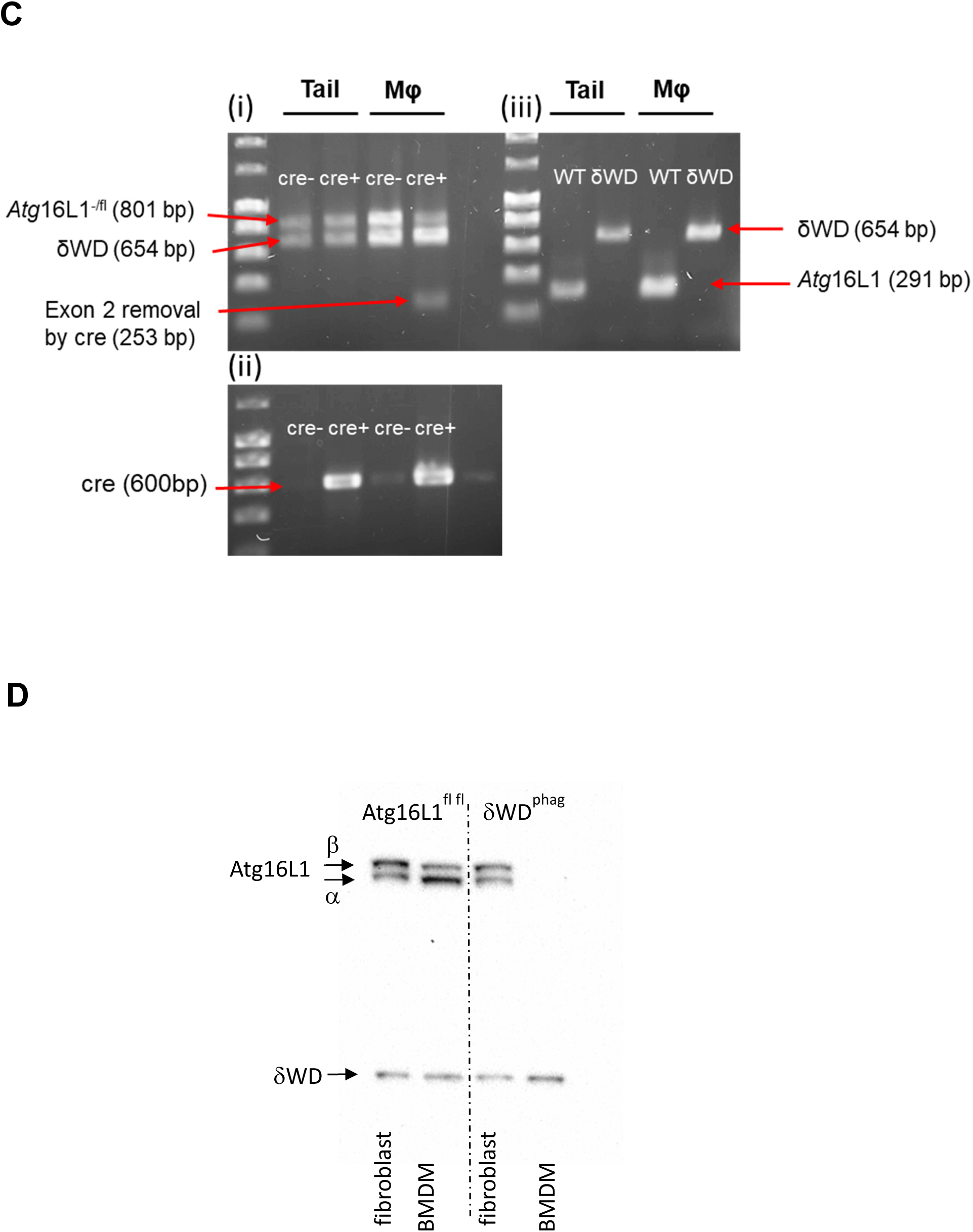
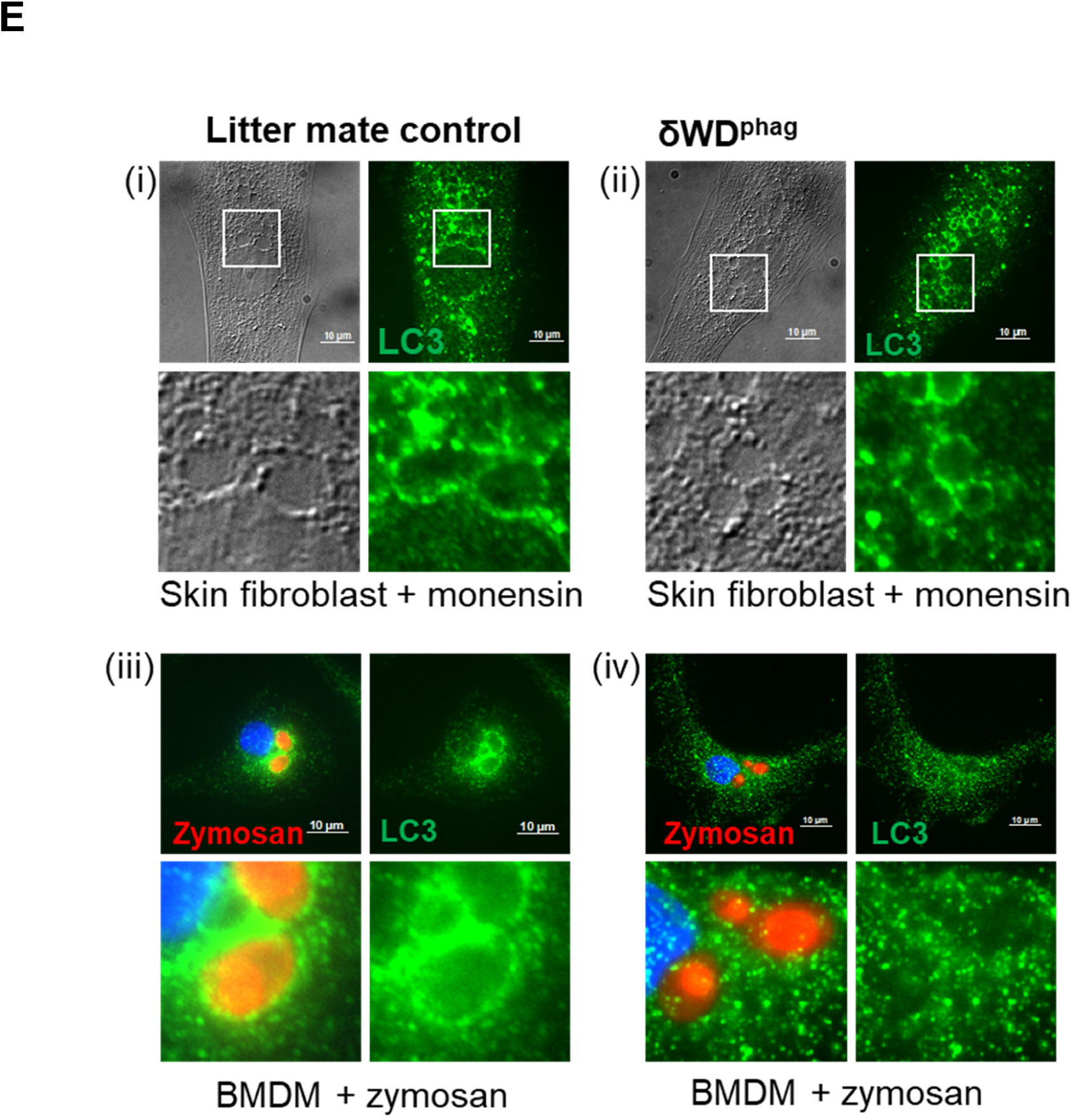
Generation of δWD^phag^ mice. **Panel A. *Breeding strategy***. Homozygous δWD mice carrying LysMcre were crossed with *Atg*16L1^fl/fl^ mice. 50% of progeny are *Atg*16L1^fl/δWD^ and carry LysMcre. Cre recombinase expressed in myeloid cells of these mice inactivates *Atg*16L1 by removing exon 2 from *Atg*16L1 (δWD^phag^). The myeloid cells only express δWD. Cre recombinase is not expressed in non-myeloid tissues and *Atg*16L1 is preserved to power autophagy. 50% of progeny provide littermate controls because they lack LysMcre and preserve *Atg*16L1 in all tissues. **Panel B. *Genome map and PCR primers for analysis of the Atg16L1 genotype***. Unmodified *Atg16*L1 is identified using primers flanking exon 2 (223, 226) and exon 6 (290 and 291). The δWD allele was generated by inserting a stop codon into exon 6 and this increases the size of the PCR product of exon 6 from 291 bp to 639 bp. In *Atg*16L1^fllfl^ loxp sites flanking exon 2 in Atg16L1 increase the PCR product of exon 2 from 654 bp to 801 bp, while removal of exon 2 by cre recombinase reduces the PCR product of exon 2 from 801 bp to 253 bp. **Panel C. *Genotyping δWD***^***phag***^ ***mice***. DNA extracted from mouse tail tissue or bone marrow derived macrophages (MF) was analysed by PCR. **(i)**. Samples from δWD^phag^ mice (indicated by cre+) and littermate controls (cre-). The 253bp PCR product seen in macrophage DNA of cre+ δWD^phag^ strains indicates specific removal of exon 2 from *Atg*16L1 in myeloid cells. **(ii)**. PCR primers verify presence of cre recombinase (cre+). iii). Genotyping of wild type and δWD strains showing predicted changes in size of PCR product from exon 6. **Panel D. Tissue specific expression of ATG16L1 and δWD**. Skin fibroblasts and bone marrow derived macrophages (BMDM) isolated from *Atg*16L1δWD^phag^ mice (δWD^phag^) and littermate controls were analysed by western blot. Skin fibrobalasts and BMDM from control mice lacking LysMcre (Atg16L1^fl/fl^) express full length 70kDa α and β isoforms of ATG16L1 and the truncated δWD at 25kDa. δWD^phag^ mice express LysMcre indicated by the removal of full length ATG16L1 from BMDM but not skin fibroblasts. **Panel E. Functional analysis of δWD**^**phag**^ **mice** ***Panels (i) and (ii). Analysis of non-canonical autophagy***/***LC3 associated endocytosis in fibroblasts from δWD***^***phag***^ ***mice***. Skin fibroblasts isolated from *Atg*16L1δWD^phag^ mice (δWD^phag^) and litter mate controls were incubated with monensin to induce LC3 associated endocytosis, fixed and immunostained for LC3. Fibroblasts from δWD^phag^ mice are able to recruit LC3 (green) to swollen endo-lysosome compartments in a similar way to those from littermate control mice. ***Panels (iii) and (iv). Analysis of non-canonical autophagy/LAP in BMDM from δWD***^***phag***^ ***mice***. BMDMs isolated from *Atg*16L1δWD^phag^ mice (δWD^phag^) and litter mate controls were incubated with zymosan for 30 min, fixed and immunostained for LC3. BMDMs from δWD^phag^ mice are unable to recruit LC3 (green) to phagosomes containing zymosan (red).

**Fig. S8.**
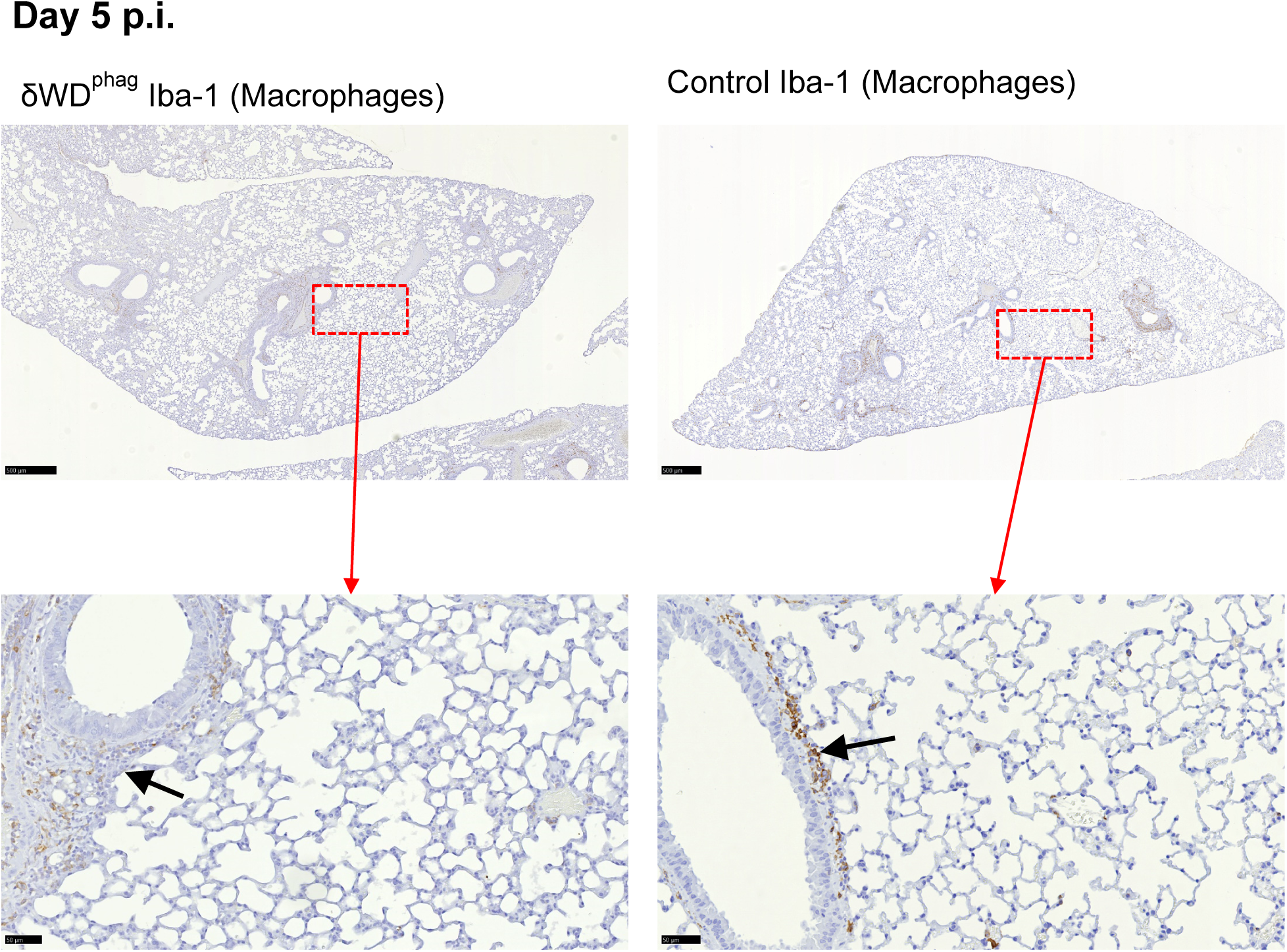
Mice deficient in non-canonical autophagy/LAP in phagocytes control IAV infection and do not show increased lung inflammation. δWD^phag^ that are deficient in non-canonical autophagy in phagocytes and littermate control mice were infected i.n. with 10^3^ pfu IAV X31. Lung tissues were harvested at 5 d p.i. Macrophages were detected by IH using anti-Iba-1, visualized with DAB and counter-stained with hematoxylin. Scale bars represent 500 µm (upper panels) and 50 µm (lower panels). Micrographs of representative areas from lungs of six mice are shown. Both δWD^phag^ and control mice show little inflammatory response in the parenchyma and mild, macrophage-rich peri-bronchiolar infiltration (black arrows).

**Fig. S9.**
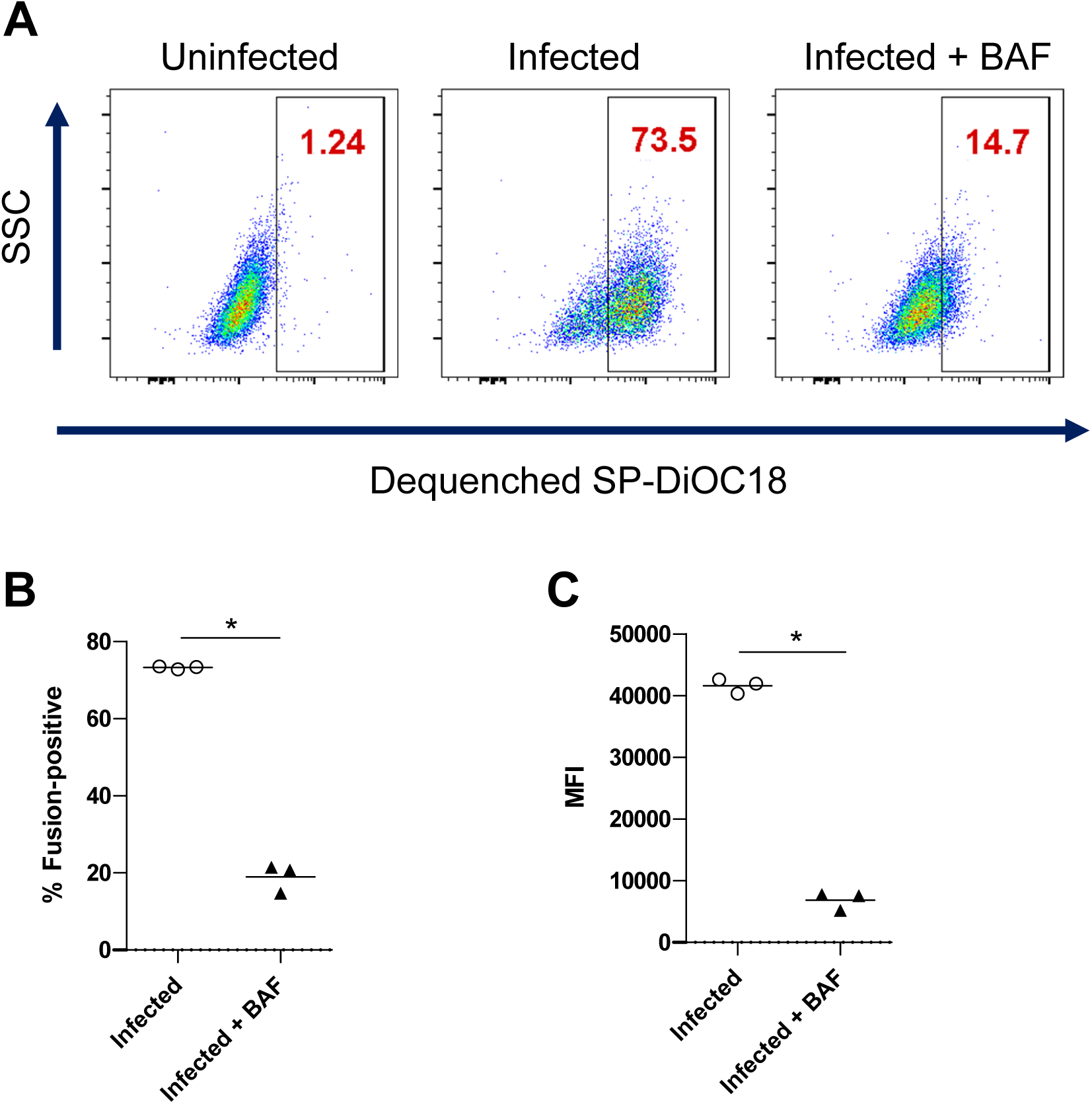
Fluorescent signal in cells infected with SP-DiOC18 labelled IAV is pH-dependent fusion. Sub-confluent monolayers of MEFs were incubated with dual-labelled (SP-DiOC18/R18) IAV at 4°C for 45 min and warmed to 37°C for 90 min. pH-dependent fusion was assessed by adding bafilomycin A1 (BAF) to some wells. Cells were harvested by trypsinisation, fixed in PFA and analysed by flow cytometry. **(A)**. Representative plots showing de-quenched SP-DiOC18. Numbers in the gates show percentage of cells positive for fusion as determined by comparison to uninfected cells. **(B)** Percentage of cells positive for fusion **(C)** Median fluorescence intensity of dequenched SP-DiOC18 signal. **B** and **C** show individual replicates with a bar at the mean and were compared using Mann-Whitney *U* test.

## Notes

### Competing Interest Statement

The authors have declared no competing interest.

### Summary of Updates

Revision includes new in vitro fusion assay data (Figure 6 and S9) that shows the WD and linker domain of ATG16L1 inhibit virus entry by affecting fusion of virus and endosome membranes. Yohei Yamauchi is also added as co-author

## REFERENCES

Akoumianaki T, Kyrmizi I, Valsecchi I, Gresnigt MS, Samonis G, Drakos E, Boumpas D, Muszkieta L, Prevost MC, Kontoyiannis DP, Chavakis T, Netea MG, van de Veerdonk FL, Brakhage AA, El-Benna J, Beauvais A, Latge JP, Chamilos G (2016) Aspergillus Cell Wall Melanin Blocks LC3-Associated Phagocytosis to Promote Pathogenicity. Cell Host Microbe 19: 79–90

Akram KM, Moyo NA, Leeming GH, Bingle L, Jasim S, Hussain S, Schorlemmer A, Kipar A, Digard P, Tripp RA, Shohet RV, Bingle CD, Stewart JP (2018) An innate defense peptide BPIFA1/SPLUNC1 restricts influenza A virus infection. Mucosal Immunol 11: 71–81

Belser JA, Gustin KM, Maines TR, Blau DM, Zaki SR, Katz JM, Tumpey TM (2011) Pathogenesis and transmission of triple-reassortant swine H1N1 influenza viruses isolated before the 2009 H1N1 pandemic. J Virol 85: 1563–72

Berryman S, Brooks E, Burman A, Hawes P, Roberts R, Netherton C, Monaghan P, Whelband M, Cottam E, Elazar Z, Jackson T, Wileman T (2012) Foot-and-mouth disease virus induces autophagosomes during cell entry via a class III phosphatidylinositol 3-kinase-independent pathway. J Virol 86: 12940–53

Boada-Romero E, Serramito-Gomez I, Sacristan MP, Boone DL, Xavier RJ, Pimentel-Muinos FX (2016) The T300A Crohn’s disease risk polymorphism impairs function of the WD40 domain of ATG16L1. Nat Commun 7: 11821

Boulo S, Akarsu H, Ruigrok RW, Baudin F (2007) Nuclear traffic of influenza virus proteins and ribonucleoprotein complexes. Virus Res 124: 12–21

Davidson S, Crotta S, McCabe TM, Wack A (2014) Pathogenic potential of interferon alphabeta in acute influenza infection. Nat Commun 5: 3864

Delgado MA, Elmaoued RA, Davis AS, Kyei G, Deretic V (2008) Toll-like receptors control autophagy. EMBO J 27: 1110–21

Dooley HC, Razi M, Polson HE, Girardin SE, Wilson MI, Tooze SA (2014) WIPI2 links LC3 conjugation with PI3P, autophagosome formation, and pathogen clearance by recruiting Atg12-5-16L1. Mol Cell 55: 238–52

Fletcher K, Ulferts R, Jacquin E, Veith T, Gammoh N, Arasteh JM, Mayer U, Carding SR, Wileman T, Beale R, Florey O (2018) The WD40 domain of ATG16L1 is required for its non-canonical role in lipidation of LC3 at single membranes. EMBO J 37

Florey O, Gammoh N, Kim SE, Jiang X, Overholtzer M (2015) V-ATPase and osmotic imbalances activate endolysosomal LC3 lipidation. Autophagy 11: 88–99

Florey O, Kim SE, Sandoval CP, Haynes CM, Overholtzer M (2011) Autophagy machinery mediates macroendocytic processing and entotic cell death by targeting single membranes. Nat Cell Biol 13: 1335–43

Gluschko A, Herb M, Wiegmann K, Krut O, Neiss WF, Utermohlen O, Kronke M, Schramm M (2018) The beta2 Integrin Mac-1 Induces Protective LC3-Associated Phagocytosis of Listeria monocytogenes. Cell Host Microbe 23: 324–337 e5

Heckmann BL, Boada-Romero E, Cunha LD, Magne J, Green DR (2017) LC3-Associated Phagocytosis and Inflammation. J Mol Biol 429: 3561–3576

Heckmann BL, Teubner BJW, Tummers B, Boada-Romero E, Harris L, Yang M, Guy CS, Zakharenko SS, Green DR (2019) LC3-Associated Endocytosis Facilitates beta-Amyloid Clearance and Mitigates Neurodegeneration in Murine Alzheimer’s Disease. Cell 178: 536–551 e14

Herold S, Becker C, Ridge KM, Budinger GR (2015) Influenza virus-induced lung injury: pathogenesis and implications for treatment. Eur Respir J 45: 1463–78

Huang J, Canadien V, Lam GY, Steinberg BE, Dinauer MC, Magalhaes MA, Glogauer M, Grinstein S, Brumell JH (2009) Activation of antibacterial autophagy by NADPH oxidases. Proc Natl Acad Sci U S A 106: 6226–31

Hubber A, Kubori T, Coban C, Matsuzawa T, Ogawa M, Kawabata T, Yoshimori T, Nagai H (2017) Bacterial secretion system skews the fate of Legionella-containing vacuoles towards LC3-associated phagocytosis. Sci Rep 7: 44795

Iwasaki A, Pillai PS (2014) Innate immunity to influenza virus infection. Nat Rev Immunol 14: 315–28

Kemball CC, Alirezaei M, Flynn CT, Wood MR, Harkins S, Kiosses WB, Whitton JL (2010) Coxsackievirus infection induces autophagy-like vesicles and megaphagosomes in pancreatic acinar cells in vivo. J Virol 84: 12110–24

Kreibich S, Emmenlauer M, Fredlund J, Ramo P, Munz C, Dehio C, Enninga J, Hardt WD (2015) Autophagy Proteins Promote Repair of Endosomal Membranes Damaged by the Salmonella Type Three Secretion System 1. Cell Host Microbe 18: 527–37

Kyrmizi I, Ferreira H, Carvalho A, Figueroa JAL, Zarmpas P, Cunha C, Akoumianaki T, Stylianou K, Deepe GS, Jr., Samonis G, Lacerda JF, Campos A, Jr., Kontoyiannis DP, Mihalopoulos N, Kwon-Chung KJ, El-Benna J, Valsecchi I, Beauvais A, Brakhage AA, Neves NM et al. (2018) Calcium sequestration by fungal melanin inhibits calcium-calmodulin signalling to prevent LC3-associated phagocytosis. Nat Microbiol 3: 791–803

Lamprinaki D, Beasy G, Zhekova A, Wittmann A, James S, Dicks J, Iwakura Y, Saijo S, Wang X, Chow CW, Roberts I, Korcsmaros T, Mayer U, Wileman T, Kawasaki N (2017) LC3-Associated Phagocytosis Is Required for Dendritic Cell Inflammatory Cytokine Response to Gut Commensal Yeast Saccharomyces cerevisiae. Front Immunol 8: 1397

Lu Q, Yokoyama CC, Williams JW, Baldridge MT, Jin X, DesRochers B, Bricker T, Wilen CB, Bagaitkar J, Loginicheva E, Sergushichev A, Kreamalmeyer D, Keller BC, Zhao Y, Kambal A, Green DR, Martinez J, Dinauer MC, Holtzman MJ, Crouch EC et al. (2016) Homeostatic Control of Innate Lung Inflammation by Vici Syndrome Gene Epg5 and Additional Autophagy Genes Promotes Influenza Pathogenesis. Cell Host Microbe 19: 102–13

Martinez J, Cunha LD, Park S, Yang M, Lu Q, Orchard R, Li QZ, Yan M, Janke L, Guy C, Linkermann A, Virgin HW, Green DR (2016) Noncanonical autophagy inhibits the autoinflammatory, lupus-like response to dying cells. Nature 533: 115–9

Martinez J, Malireddi RK, Lu Q, Cunha LD, Pelletier S, Gingras S, Orchard R, Guan JL, Tan H, Peng J, Kanneganti TD, Virgin HW, Green DR (2015) Molecular characterization of LC3-associated phagocytosis reveals distinct roles for Rubicon, NOX2 and autophagy proteins. Nat Cell Biol 17: 893–906

Matte C, Casgrain PA, Seguin O, Moradin N, Hong WJ, Descoteaux A (2016) Leishmania major Promastigotes Evade LC3-Associated Phagocytosis through the Action of GP63. PLoS Pathog 12: e1005690

Rai S, Arasteh M, Jefferson M, Pearson T, Wang Y, Zhang W, Bicsak B, Divekar D, Powell PP, Naumann R, Beraza N, Carding SR, Florey O, Mayer U, Wileman T (2019) The ATG5-binding and coiled coil domains of ATG16L1 maintain autophagy and tissue homeostasis in mice independently of the WD domain required for LC3-associated phagocytosis. Autophagy 15: 599–612

Ramos I, Fernandez-Sesma A (2015) Modulating the Innate Immune Response to Influenza A Virus: Potential Therapeutic Use of Anti-Inflammatory Drugs. Front Immunol 6: 361

Roberts R, Al-Jamal WT, Whelband M, Thomas P, Jefferson M, van den Bossche J, Powell PP, Kostarelos K, Wileman T (2013) Autophagy and formation of tubulovesicular autophagosomes provide a barrier against nonviral gene delivery. Autophagy 9: 667–82

Saitoh T, Fujita N, Jang MH, Uematsu S, Yang BG, Satoh T, Omori H, Noda T, Yamamoto N, Komatsu M, Tanaka K, Kawai T, Tsujimura T, Takeuchi O, Yoshimori T, Akira S (2008) Loss of the autophagy protein Atg16L1 enhances endotoxin-induced IL-1beta production. Nature 456: 264–8

Sanjuan MA, Dillon CP, Tait SW, Moshiach S, Dorsey F, Connell S, Komatsu M, Tanaka K, Cleveland JL, Withoff S, Green DR (2007) Toll-like receptor signalling in macrophages links the autophagy pathway to phagocytosis. Nature 450: 1253–7

Skehel JJ, Wiley DC (2000) Receptor binding and membrane fusion in virus entry: the influenza hemagglutinin. Annu Rev Biochem 69: 531–69

Szretter KJ, Gangappa S, Lu X, Smith C, Shieh WJ, Zaki SR, Sambhara S, Tumpey TM, Katz JM (2007) Role of host cytokine responses in the pathogenesis of avian H5N1 influenza viruses in mice. J Virol 81: 2736–44

Tan JMJ, Mellouk N, Osborne SE, Ammendolia DA, Dyer DN, Li R, Brunen D, van Rijn JM, Huang J, Czuczman MA, Cemma MA, Won AM, Yip CM, Xavier RJ, MacDuff DA, Reggiori F, Debnath J, Yoshimori T, Kim PK, Fairn GD et al. (2018) An ATG16L1-dependent pathway promotes plasma membrane repair and limits Listeria monocytogenes cell-to-cell spread. Nat Microbiol 3: 1472–1485

Teijaro JR, Walsh KB, Rice S, Rosen H, Oldstone MB (2014) Mapping the innate signaling cascade essential for cytokine storm during influenza virus infection. Proc Natl Acad Sci U S A 111: 3799–804

Ullrich S, Munch A, Neumann S, Kremmer E, Tatzelt J, Lichtenthaler SF (2010) The novel membrane protein TMEM59 modulates complex glycosylation, cell surface expression, and secretion of the amyloid precursor protein. J Biol Chem 285: 20664–74

Wharton SA, Belshe RB, Skehel JJ, Hay AJ (1994) Role of virion M2 protein in influenza virus uncoating: specific reduction in the rate of membrane fusion between virus and liposomes by amantadine. J Gen Virol 75 (Pt 4): 945–8

Yamayoshi S, Kawaoka Y (2019) Current and future influenza vaccines. Nat Med 25: 212–220

Yang CS, Shin DM, Kim KH, Lee ZW, Lee CH, Park SG, Bae YS, Jo EK (2009) NADPH oxidase 2 interaction with TLR2 is required for efficient innate immune responses to mycobacteria via cathelicidin expression. J Immunol 182: 3696–705

